# The global phylogenetic landscape and nosocomial spread of the multidrug-resistant opportunist *Stenotrophomonas maltophilia*

**DOI:** 10.1101/748954

**Authors:** Matthias I Gröschel, Conor J Meehan, Ivan Barilar, Margo Diricks, Aitor Gonzaga, Matthias Steglich, Oscar Conchillo-Solé, Isabell-Christin Scherer, Uwe Mamat, Christian F. Luz, Katrien De Bruyne, Christian Utpatel, Daniel Yero, Isidre Gibert, Xavier Daura, Stefanie Kampmeier, Nurdyana Abdul Rahman, Michael Kresken, Tjip S van der Werf, Ifey Alio, Wolfgang R. Streit, Kai Zhou, Thomas Schwartz, John W A Rossen, Maha R Farhat, Ulrich E Schaible, Ulrich Nübel, Jan Rupp, Joerg Steinmann, Stefan Niemann, Thomas A Kohl

**Author notes:** These authors jointly supervised the work: Jan Rupp, Joerg Steinmann, Stefan Niemann, Thomas A Kohl. Correspondence to: Prof Stefan Niemann.

## Abstract

Recent studies portend a rising global spread and adaptation of human- or healthcare-associated pathogens. Here, we analysed an international collection of the emerging, multidrug-resistant, opportunistic pathogen *Stenotrophomonas maltophilia* from 22 countries to infer population structure and clonality at a global level. We show that the *S. maltophilia* complex is divided into 23 monophyletic lineages, most of which harboured strains of all degrees of human virulence. Lineage Sm6 comprised the highest rate of human-associated strains, linked to key virulence and resistance genes. Transmission analysis identified potential outbreak events of genetically closely related strains isolated within days or weeks in the same hospitals.

**One Sentence Summary:** The *S. maltophilia* complex comprises genetically diverse, globally distributed lineages with evidence for intra-hospital transmission.

## Main Text

Recently, local transmission and global spread of hospital-acquired pathogens such as *Mycobacterium abscessus* and *Mycobacterium chimaera* were revealed by whole genome sequencing (WGS), thereby challenging the prevailing concepts of disease acquisition and transmission of these pathogens in the hospital setting^1–3^. Global genome-based collections are missing for other emerging pathogens such as *Stenotrophomonas maltophilia*, listed by the World Health Organization as one of the leading drug-resistant nosocomial pathogens worldwide^4^*. S. maltophilia* is ubiquitously found in natural ecosystems and is of importance in environmental remediation and industry^5, 6^. *S. maltophilia* is an important cause of hospital-acquired drug-resistant infections with a significant attributable mortality rate in immunocompromised patients of up to 37.5%^7–9^. Patients under immunosuppressive treatment and those with malignancy or pre-existing inflammatory lung diseases such as cystic fibrosis are at particular risk of *S. maltophilia* infection^10^. Although almost any organ can be affected, mere colonization needs to be discriminated from infections that mainly manifest as respiratory tract infections, bacteraemia, or catheter-related bloodstream infections^5^. Yet, the bacterium is also commonly isolated from wounds and, in lower frequency, in implant-associated infections^11, 12^. Furthermore, community acquired infections have also been described^13^. Treatment options are limited by resistance to a number of antimicrobial classes such as most ß-lactam antibiotics, cephalosporins, aminoglycosides, and macrolides through the intrinsic resistome, genetic material acquired by horizontal transfer, as well as non-heritable adaptive mechanisms^14, 15^.

To date, no large-scale genome-based studies on the population structure and clonality of *S. maltophilia* in relation to human disease have been conducted. Previous work indicated the presence of at least 13 lineages or species-like lineages in the *S. maltophilia* complex, defined as *S. maltophilia* strains with 16S rRNA gene sequence similarities > 99.0%, with nine of these potentially human-associated^8, 16–19^. These *S. maltophilia* complex lineages are further divided into four more distantly related lineages (Sgn1-4) and several *S. maltophilia sensu lato* and *sensu stricto* lineages^16, 20^. The *S. maltophilia* type strain K279a, isolated from a patient with bloodstream infection, serves as an indicator strain of the lineage *S. maltophilia sensu stricto*^20^.

To understand the global population structure of the *S. maltophilia* complex and the potential for global and local spread of strains, in particular of human-associated lineages, we performed a large-scale genome-based phylogenetic and cluster analysis of a global collection of newly sequenced *S. maltophilia* strains together with publicly available whole genome data.

## Results

### Strain collection and gene-by-gene analysis

To allow for standardised WGS-based genotyping and gene-by-gene analysis of our dataset, we first created an *S. maltophilia* complex whole genome multilocus sequence typing (wgMLST) scheme. This approach, implemented as core genome MLST, has been widely used in tracing outbreaks and transmission events for a variety of bacterial species^21–23^. The use of a wgMLST scheme allows to analyze sequenced strains by their core and accessory genome^24^. Using 171 publicly available assembled genomes of the *S. maltophilia* complex that represent its currently known diversity (Data S1) we constructed a wgMLST scheme consisting of 17,603 loci (Data S2). To ensure compatibility with traditional MLST / *gyrB* typing methods, the wgMLST scheme includes the partial sequences of the seven genes used in traditional MLST as well as the *gyrB* gene^25^ (Table S1).

**Table 1.** Site and date of isolation for the strains comprising the four d10 clusters isolated from the same geographic location within at most an eight weeks time span.

| Strain | Lineage | Cluster | Isolation date | Isolation place | Clinical source |
| --- | --- | --- | --- | --- | --- |
| PEG-257 | Sm2 | 24 | October 4th, 2013 | Hospital A | respiratory tract |
| PEG-258 |  |  | October 7th, 2013 |  | wound swap |
| PEG-263 |  |  | October 9th, 2013 |  | respiratory tract |
| PEG-266 |  |  | October 14th, 2013 |  | respiratory tract |
| PEG-267 |  |  | October 14th, 2013 |  | respiratory tract |
| PEG-268 |  |  | October 14th, 2013 |  | sputum |
| PEG-328 | Sm18 | 26 | October 21st, 2013 | Hospital B | respiratory tract |
| PEG-329 |  |  | October 21st, 2013 |  |  |
| PEG-331 |  |  | December 9th, 2013 |  |  |
| PEG-351 | Sm13 | 27 | January 11th, 2014 | Hospital C | sputum |
| PEG-349 |  |  | January 21st, 2014 |  | respiratory tract |
| PEG-350 |  |  | January 24th, 2014 |  | respiratory tract |
| PEG-353 |  |  | January 27th, 2014 |  | respiratory tract |
| 943974Y | Sm12 | 31 | January 28th, 2014 | Hospital D | endoscope |
| 944632W |  |  | February 17th, 2014 |  |  |
| 944796D |  |  | February 21st, 2014 |  |  |
| 945570W |  |  | March 24th, 2014 |  |  |

To investigate the global phylogeographic distribution of *S. maltophilia*, we gathered WGS data of 2,389 strains from 22 countries and four continents, which were either collected and sequenced in this study or had sequence data available in public repositories (Data S3). All genome assemblies of the study collection passing quality thresholds (Fig. S1, Data S4) were analyzed with the newly created wgMLST scheme. Upon duplicate removal, filtering for sequence quality, and removal of strains with fewer than 2,000 allele calls in the wgMLST scheme, our study collection comprised 1,305 assembled genomes of majority clinical origin (87%) of which 234 were from public repositories and 1,071 newly sequenced strains. Most strains came from Germany (932 strains), the United States (92 strains), Australia (56 strains), Switzerland (49 strains), and Spain (42 strains) (Fig 1D, Data S3). WgMLST analysis resulted in an average of 4,174 (range 3,024 – 4,536) loci recovered per strain. Across the 1305 strains, most loci, 13,002 of 17,603, were assigned fewer than 50 different alleles, consistent with a large accessory genome (Fig. S2). This was confirmed by calculation of the sample pan genome that yielded 17,479 loci, with 2,844 loci (16.3%) present in 95% and 1,274 loci (7.3%) present in 99% of strains (Fig. S3). The genome sizes ranged from 4.04 Mb to 5.2 Mb.

**Fig. 1.**
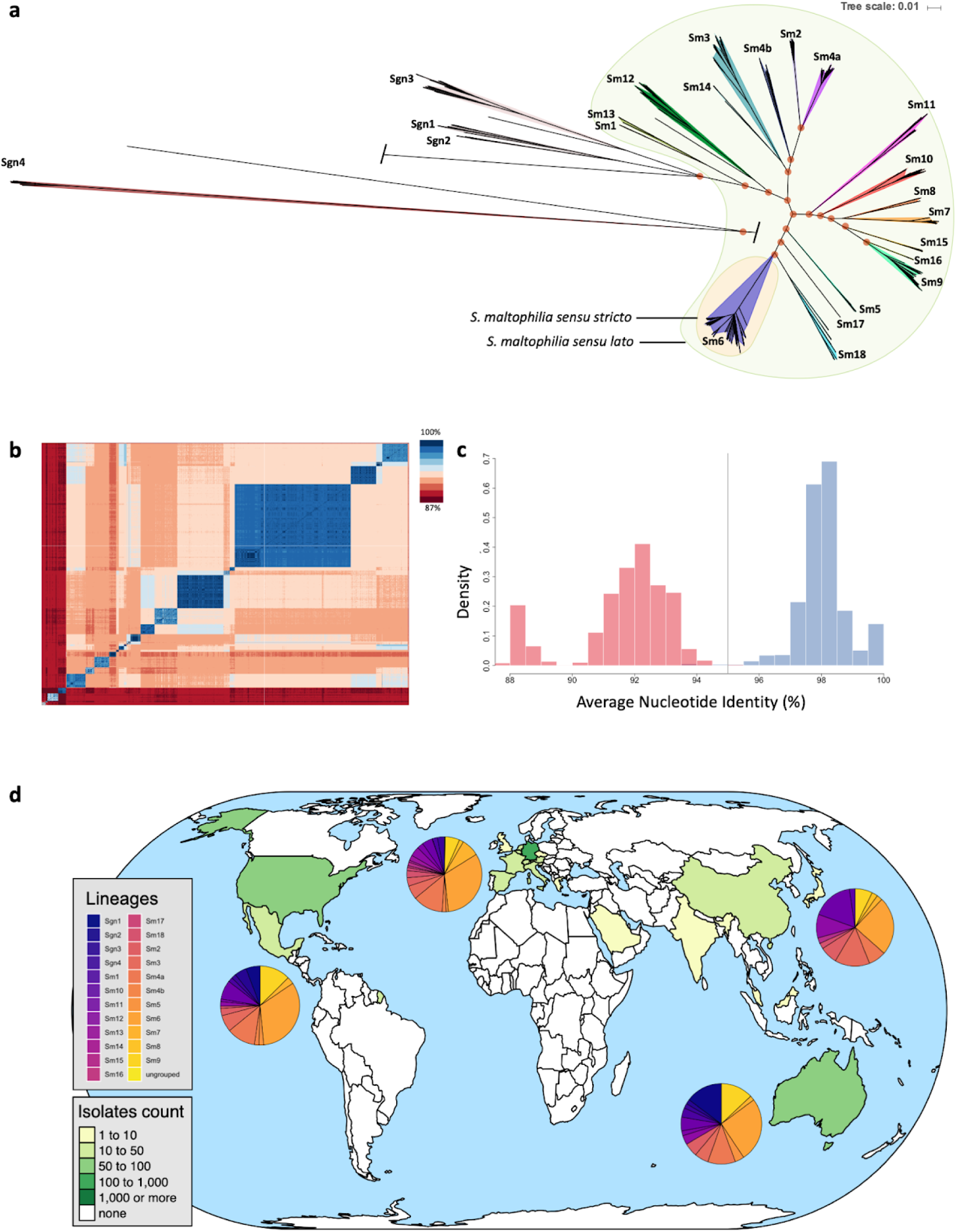
The global population structure of the *S. maltophilia* complex is composed of 23 monophyletic, globally distributed lineages. **a,** Unrooted maximum likelihood phylogenetic tree of 1,305 *S. maltophilia* strains displaying the known population diversity of the *S. maltophilia* complex. The tree was built using RAxML on the sequences of 1,274 concatenated core genome genes. Groups as defined by hierarchical Bayesian clustering are marked with shaded colours, and group numbers are indicated at the tree leafs of each corresponding group. orange shading = *S. maltophilia sensu stricto*; green shading = *S. maltophilia sensu lato*; 100% support values for the main branches are indicated with red circles. **b,** Pairwise Average Nucleotide Identity comparison calculated for 1,305 *S. maltophilia* strains shown on a heatmap with blue indicating high and red indicating low nucleotide identity. **c,** Histogram of pairwise average nucleotide identity (ANI) values, illustrating that strains of the same lineage are highly similar at the nucleotide level with ANI values above 95% (depicted in blue). Inter-lineage comparisons (in red colour) reveal low genetic identity between strains. The currently accepted species delimitation threshold at 95% is shown as a grey vertical line. **d,** Geographic origin of the 1,305 *S. maltophilia* strains comprising the study collection indicated on a global map. The green/yellow colour code indicates the number of strains obtained per country. The distribution of phylogenetic lineages per continent is displayed as colour-coded pie charts.

### Delineation of the *S. maltophilia* complex within the *Stenotrophomonas* genus

To obtain insights on the relatedness of the members of the *Stenotrophomonas* genus we first analyzed all *Stenotrophomonas* species with available sequence data using the wgMLST scheme. We recovered between 380 loci in *S. dokdonensis* to a maximum of 1,677 in *S. rhizophila*, with *S. terrae, S. panacihumi, S. humi, S. chelatiphaga, S. daejeonensis, S. ginsengisoli*, *S. koreensis*, and *S. acidaminiphilia* species obtaining. For the strain *S. maltophilia* JCM9942 (Genbank accession GCA_001431585.1), only 982 loci were detected. Interestingly, the 16S rRNA gene sequence of JCM9942 matched to *S. acidaminiphila*, and the JCM9942 16S rRNA sequence is only 97.3% identical with that of *S. maltophilia* K279a. In contrast, the number of recovered loci matches those of *S. maltophilia* strains for *P. geniculata* (4,060 loci), *S. lactitubi* (3,805 loci), *S. pavanii* (3,623 loci), and *S. indicatrix* (3,678 loci). Here, 16S rRNA sequence comparison to *S. maltophilia* K279a would support the inclusion of the first 3 of the aforementioned species with 16S rRNA sequence identity of >99.1% into the *S. maltophilia* complex^19, 26^.

### *S. maltophilia* complex comprises 23 distinct monophyletic lineages

To investigate the global diversity of the *S. maltophilia* complex, a maximum likelihood phylogeny was inferred from a concatenated sequence alignment of the 1,274 core loci present in 99% of the 1,305 *S. maltophilia* strains of our study collection (Fig. 1A). Hierarchical Bayesian analysis of population structure (BAPS), derived from the core single nucleotide polymorphism (SNP) results, clustered the 1,305 genomes into 23 monophyletic lineages named Sgn1-Sgn4 and Sm1-Sm18, comprising 17 previously suggested and six hitherto unknown lineages (Sm13 - Sm18). For consistency we used and amended the naming convention of lineages from previous reports^16, 18^. In concordance with these studies^16, 18^, we found a clear separation of the more distantly related lineages Sgn1-Sgn4 and a branch formed by lineages Sm1-Sm18 (previously termed *S. maltophilia sensu lato*), with the largest lineage Sm6 (also known as *S. maltophilia sensu stricto*) containing most strains (n = 413) and the clinical reference strain K279a. Contrary to previous analyses, Sgn4 is the lineage most distantly related to the rest of the strains^16^. The division into the 23 lineages is also clearly supported by an Average Nucleotide Identity (ANI) analysis (Fig. 1B and Fig. 1C). ANI comparisons of strains belonging to the same lineage was above 95%, and comparisons of strains between lineages were below 95%.

To evaluate structural genomic variation across the various lineages, we compiled a set of 20 completely closed genomes covering the 15 major phylogenetic lineages of both environmental and human-invasive or human-non-invasive isolation source. These genomes were either procured from the NCBI (n = 8) or newly sequenced on the PacBio platform (n = 12) (Table S2). Interestingly, no plasmids were detected in any of the genomes. A genome-wide alignment of the 20 genomes including the K279a reference strain demonstrated considerable variation in both structure and size between strains of different lineages and even strains of the same lineage (Fig. S4). Several phage-related, integrative and conjugative mobile elements were observed across the genomes.

### Phylogenetic lineages are globally distributed and differ in their human association

We next analyzed the global distribution of strains of the lineages defined above and found that 8 (Sm2, Sm3, Sm4a, Sm6, Sm7, Sm9, Sm10, and Sm12) are represented on all continents sampled within this study, with strains of lineage Sm6 accounting for the largest number of strains globally and the largest proportion on each sampled continent (Fig. 1D, Fig. 2A). To further investigate whether the lineages correlate with isolation source, particularly with regard to human host adaptation, we classified the isolation source of the *S. maltophilia* strains into five categories. Strains were considered environmental (n = 117) if found in natural environments e.g. in the rhizosphere, and anthropogenic if swabbed in human surroundings such as patient room sink or sewage (n = 52). Human-invasive (n = 133) was used for isolates from blood, urine, drainage fluids, biopsies, or in cerebrospinal fluid, human-non-invasive (n = 353) refers to colonizing isolates from swabs of the skin, perineum, nose, oropharynx, wounds as well as intravascular catheters, and human-respiratory (n = 524) includes strains from the lower respiratory tract below the glottis and sputum collected from cystic fibrosis patients. For 126 strains, no information on their isolation source was available and, thus, these were not included in this analysis.

**Fig. 2.**
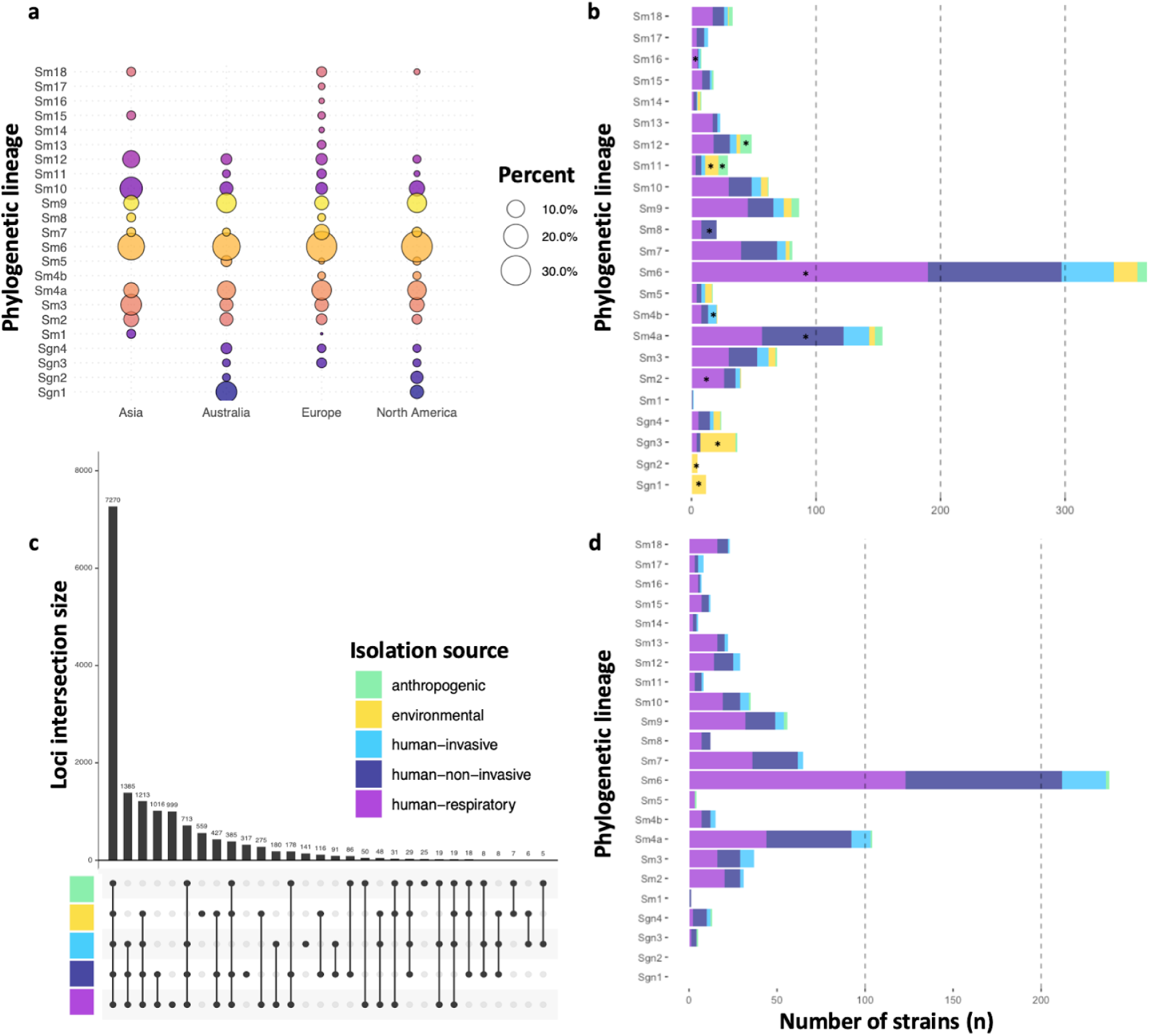
Global distribution of lineages, their composition by isolation source and contribution to phylogenetic lineage total number of strains. **a,** Bubble plot illustrating the proportion of lineages per continent. **b,** Barplot showing the number of strains per lineage coloured by isolation source for the entire strain collection. **c,** Intersection plot showing the relationship of 1,179 *S. maltophilia* strains stratified by isolation source. The plot visualizes intersections and their aggregates, illustrating that the largest number of loci is shared by the 1,179 strains, the second largest group of loci is shared by the strains of human source (invasive, non-invasive, respiratory), and so on. **d,** Barplot for the prospectively sampled representative collection of human-associated *S. maltophilia* complex strains per lineage coloured by human-invasive, human-non-invasive, or human-respiratory. * < .05 Fisher’s exact test (for n < 5) or test of equal and given proportions.

The more distantly related lineages Sgn1 (85%), Sgn2 (83%), Sgn3 (74%), and also Sm11 (34%) contained significantly more environmental strains (p < .001, test of equal or given proportions or Fisher’s exact test for n < 5), whereas strains of lineages Sm4a and Sm6 (2% and 5%, p < .001) were minority environmental (Fig. 2B and 2D, Fig. 4, Table S3). Anthropogenic strains were found at higher proportions in lineages Sm11 and Sm12 (22% and 17%, p < .001). Strains of lineage Sm4b were likely to be classified as human-invasive (32%, p = .02), and strains of lineage Sm4a and Sm8 were more likely to be human-non-invasive (40%, p < .001 and 57%, p = .03, respectively). Sgn3 contained only few human-non-invasive strains (8%, p = 0.02). Strains of lineages Sm6 (22%, p < .001), Sm2 (27%, p = .04) and Sm 13 (52%, p = .03) were linked to the human-respiratory isolation source. Strains of Sgn3 (5%, p < .001) and Sm11 (3%, p < .001) were less likely to be isolated from the human respiratory tract (Fig. 2B).

The majority of strains sequenced within this study were prospectively collected through a hospital consortium across Germany, Austria, and Switzerland (n = 741) (Fig. 2D). Restricting to this collection of human-associated strains from hospitalised patients, the most common lineage was Sm6 (33%) and lineages Sgn1-3 and Sm11, found to be environmentally-associated in public data, were either not present or represented a minor proportion (< 0.1%) of the collection.

### Genomic features of human-associated and environmental *S. maltophilia* lineages

To better understand *S. maltophilia’*s genetic adaptation to human vs environmental niches we examined genes unique to strains isolates from different sources. Human-associated strains harboured unique loci not shared by environmental, or anthropogenic strains. Human associated strains as a group shared a larger pool of unique loci (1,385) than shared with strains isolated from an environmental source (1,213) (Fig. 2C). Among the three human-associated categories, human-respiratory strains possessed the largest number of unique loci (999). Overall, we found 6,836 loci being present uniquely in human-associated strains and 932 only in environmental strains (Table S4). We next queried for lineage-specific sets of loci, i.e. present wgMLST loci that are unique to the lineage and shared among all members of that particular lineage. The *S. maltophilia sensu lato* lineages, Sm1 - Sm18, as a group, harboured 9,327 unique loci, while the more distantly related lineages, Sgn1 - 4, contained 779 unique loci. Lineage Sm6 (963 loci), Sgn3 (727 loci) and Sm3 (257 loci) displayed the highest number of lineage-specific loci. Six lineages with only few strains exhibited no unique loci (Sgn1, Sgn4, Sm1, Sm13, Sm14, and Sm16).

### Resistome and virulence characteristics of *S. maltophilia*

We next screened our collection to detect potential resistance genes, e.g. chromosomally-encoded antibiotic resistance genes including efflux pumps^7, 20, 29^. We could identify members of the five major families of efflux transporters with high frequency in our strain collection (Fig 3A)^20, 30^. Aminoglycoside modifying enzymes were encoded in 6.1% of strains (aminoglycoside-acetyltransferases) and 66% of strains (aminoglycoside-phosphotransferases), respectively, with five strains also harbouring aminoglycoside-nucleotidyltransferases. We observed that these enzyme families were unequally distributed among lineages, which preferentially contained either of the two major types. Taken together, 69% of the strains of our collection featured aminoglycoside-modifying enzymes. Other enzymes implicated in aminoglycoside resistance are the proteases ClpA and HtpX that were present in 96.9% and 98.8% of the strains investigated, respectively^31^. The *S. maltophilia* K279a genome encodes two ß-lactamases, the metallo-ß-lactamase *blaL1* and the inducible Ambler class A ß-lactamase *blaL2*^32^. While *blaL1* was found in 83.2% of our strains, *blaL2* was detected in only 63.2%. Interestingly, strains of some lineages lacked the *blaL2* gene, i.e. Sgn4, Sm1, Sm12, Sm13, and Sm16. Sm4a was the only lineage where no *blaL1* was found. Only one isolate encoded the oxacillin hydrolyzing class D ß-lactamase OXA. We noted a few strains harbouring the Type B chloramphenicol-*O*-acetyltransferase CatB (0.6%). The sulfonamide resistance-conferring *sul1* was seen in 17 strains (1.3%), and *sul2* was found in only five strains (0.4%), mostly occurring in human-associated or anthropogenic strains. This hints towards a low number of trimethoprim/sulfamethoxazole resistant strains in our collection, which is the recommended first-line agent for the treatment of *S. maltophilia* infection^14^.

**Fig. 3.**
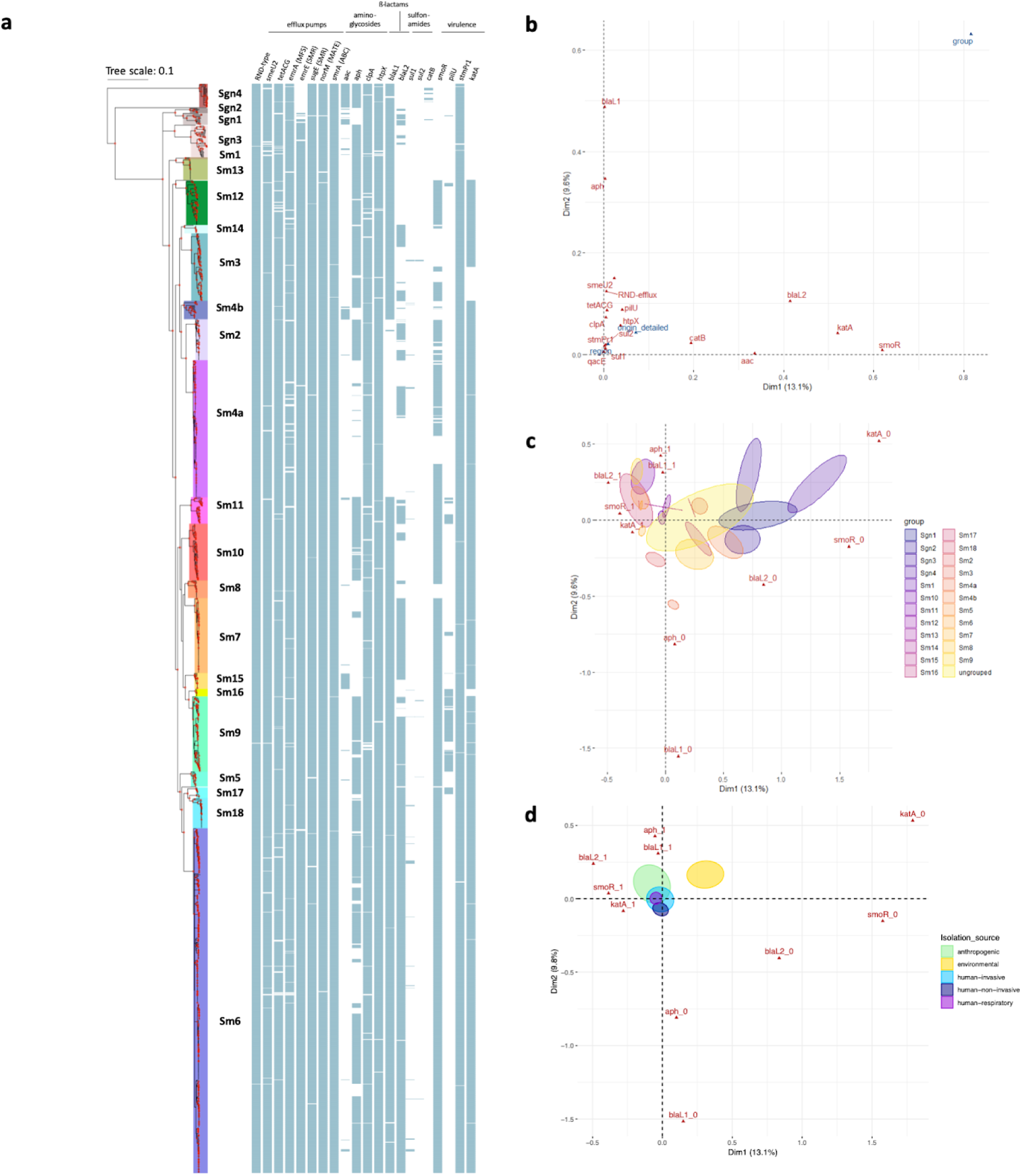
Resistance and virulence gene analysis. **a,** Maximum likelihood phylogenetic tree based on 1,274 core gene sequences of 1,305 *S. maltophilia* complex strains. The coloured shading of the lineages represents the groups found by Bayesian clustering, with lineages names given. 100% branch support is indicated by red dots. The pattern of gene presence (blue coloured line) or absence (white) is displayed in columns next to the tree, showing, from left to right, selected efflux pump genes: resistance-nodulation-cell-division (RND)-type efflux pumps, *smeU2* as part of the five-gene RND efflux pump operon *smeU1-V-W-U2-X*, *tatACG*, *emrA* of the Major Facilitator Superfamily (MFS), *emrE* and *sugE* of the small-multidrug-resistance (SMR) efflux pump family, *norM* of the MATE family, and *smrA* of the ABC-type family; the aminoglycoside acetyltransferase *aac* and phosphotransferase *aph*, *clpA*, *htpX*, the ß-lactamases *blaL1* and *blaL2*, the *sul1* and *sul2* genes encoding dihydropteroate synthases, *catB*, and the virulence genes *smoR*, *pilU*, *stmPr1*, and *katA*. **b,** Variable correlation plot visualising the 17 active variables in red and three supplementary variables in blue. **c,** Factor individual biplot map of phylogenetic lineages, indicated by their 99% confidence intervals (ellipses) across the first two MCA dimensions. The five highest contributing active variables are shown in red with 0 denoting absence and 1 presence of this variable; **d,** Factor individual biplot map of the isolation source.

We investigated the presence of virulence genes in our collection. SmoR is involved in quorum sensing and swarming motility of *S. maltophilia* and was observed in 80% of our strains^33^. PilU, a nucleotide-binding protein that contributes to Type IV pilus function, was found in 9% of strains and mainly in lineages Sm9 and Sm11^34^. StmPr1 is a major extracellular protease of *S. maltophilia* and is present in 99.2% of strains^35^. KatA is a catalase mediating increased levels of persistence to hydrogen peroxide-based disinfectants and was found in 86.6% of strains^36^. Taken together, *S. maltophilia* strains harbour a number of resistance-conferring as well as virulence genes, some of which are unequally distributed over the lineages.

To further investigate the correlation of the resistance and virulence profiles of the strains with geographic origin, isolation source, and phylogenetic lineage, we used Multiple Correspondence Analysis (MCA) to analyze potential associations. A total of 17 genes, derived from virulence databases^37, 38^, that were not present in all, or in a minority of strains were selected to serve as active variables for the MCA. As expected with a complex dataset, the total variance explained by the MCA model was relatively low (Fig. S5A). Nevertheless, from examining the first two dimensions of the MCA we noted that the genes *smoR*, *katA*, *blaL2*, *aac,* and *catB* correlate with the first dimension of the MCA, while genes *blaL1*, *aph*, *smeU2*, and genes encoding RND-efflux pumps are corresponding to the second dimension (Fig. 3B, Fig. S5B). When introducing geographic origin, isolation source, and phylogenetic lineages as supplementary variables to the model, we observed a strong correlation of phylogenetic lineages with both dimensions, while little to no correlation was observed for isolation source and geographic origin (Fig. 3B). This indicates that virulence and resistance profiles of the 17 genes are largely lineage-specific, with little impact of geographic origin or isolation source. However, we found a clear separation of the environmental strains from the rest of the collection when analyzing the impact of human versus environmental habitat on the observed variance (Fig. 3D). A more detailed analysis of the observed lineage-specific variation reveals that the more distantly related lineages Sgn1-4 are characterized by the lack of *smoR*, *katA,* and *blaL2* virulence and resistance determinants, whereas the human-associated lineages Sm6, Sm9, and Sm11 are strongly associated with the presence of *blaL2*, *aph*, *blaL1*, *smoR*, and *katA* (Fig. 3C). In summary, the human-associated lineages are characterized by the presence of key resistance-conferring genes.

### International presence of clonal complexes and possible local spread derived from genetic diversity analysis

The identification of widely-spread clonal complexes or potential outbreak events of *S. maltophilia* complex strains would have significant implications for preventive measures and infection control of *S. maltophilia* in clinical settings. We assessed our strain collection for circulating variants and clustered strains using the 1,274 core genome MLST loci, that were also used for phylogenetic inference, and thresholds of 100 (d100 clusters) and 10 mismatched alleles (d10 clusters) for single linkage clustering (Fig. 4A). These thresholds were chosen based on the distribution of allelic mismatches (Fig. 4B). We found 765 (63%) strains to group into 82 clusters (median cluster size 6, IQR 6 - 11.7) within 100 alleles difference. A total of 269 (21%) strains were grouped into 62 clusters within 10 alleles difference (median cluster size 4, IQR 3 - 4.7). The maximum number of strains per cluster were 45 and 12 for the d100 and d10 clusters, respectively. Interestingly, strains within d100 clonal complexes originated from different countries or cities (Fig. 5A).

**Fig. 4.**
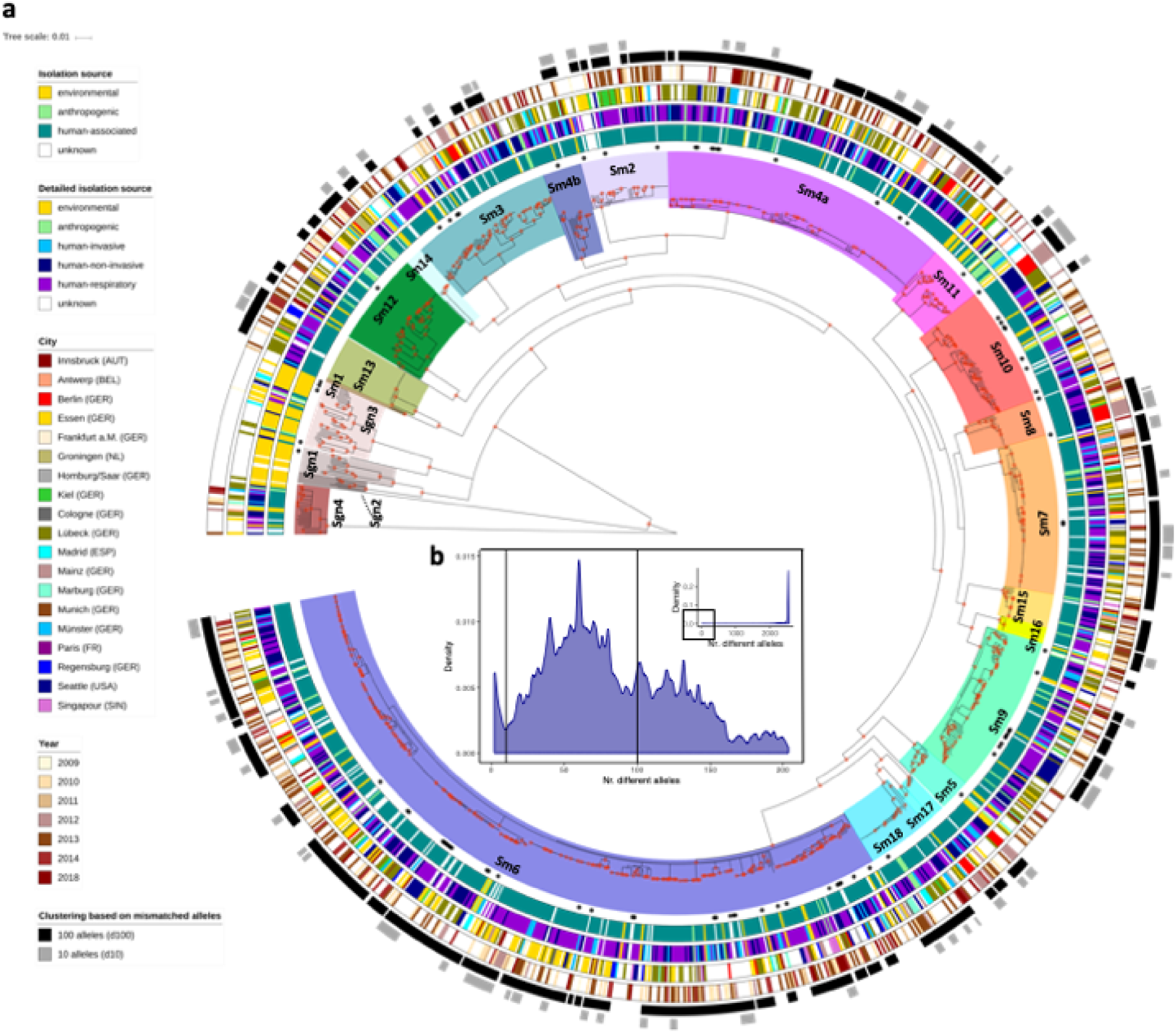
Spatiotemporal cluster analysis of 1,305 *S. maltophilia* complex strains. **a,** The coloured ranges across the outer nodes and branches indicate the 23 lineages. The black dots indicate the location of the genome datasets used for wgMLST scheme generation. The rings, from inside towards outside denote: i) the isolation source of the strains classified as either environmental, anthropogenic, human, or unknown; ii) the detailed isolation source of strains similar to the first ring with the human strains subclassified into human-invasive, human-non-invasive, and human-respiratory; iii) the city of isolation; iv) the year of isolation (where available), with light colours representing earlier years and darker brown colours more recent isolation dates. The outer rings in black-to-grey indicate the single linkage-derived clusters based on the number of allelic differences between any two strains for 100 (d100 clusters) and 10 (d10 clusters) allelic mismatches. Red dots on the nodes indicate support values of 100%. **b,** Distribution of the number of wgMLST allelic differences between pairs of strains among the 1,305 *S. maltophilia* strains. The main figure shows the frequencies of up to 200 allelic differences, while the inset displays frequencies of all allelic mismatches.

**Fig. 5.**
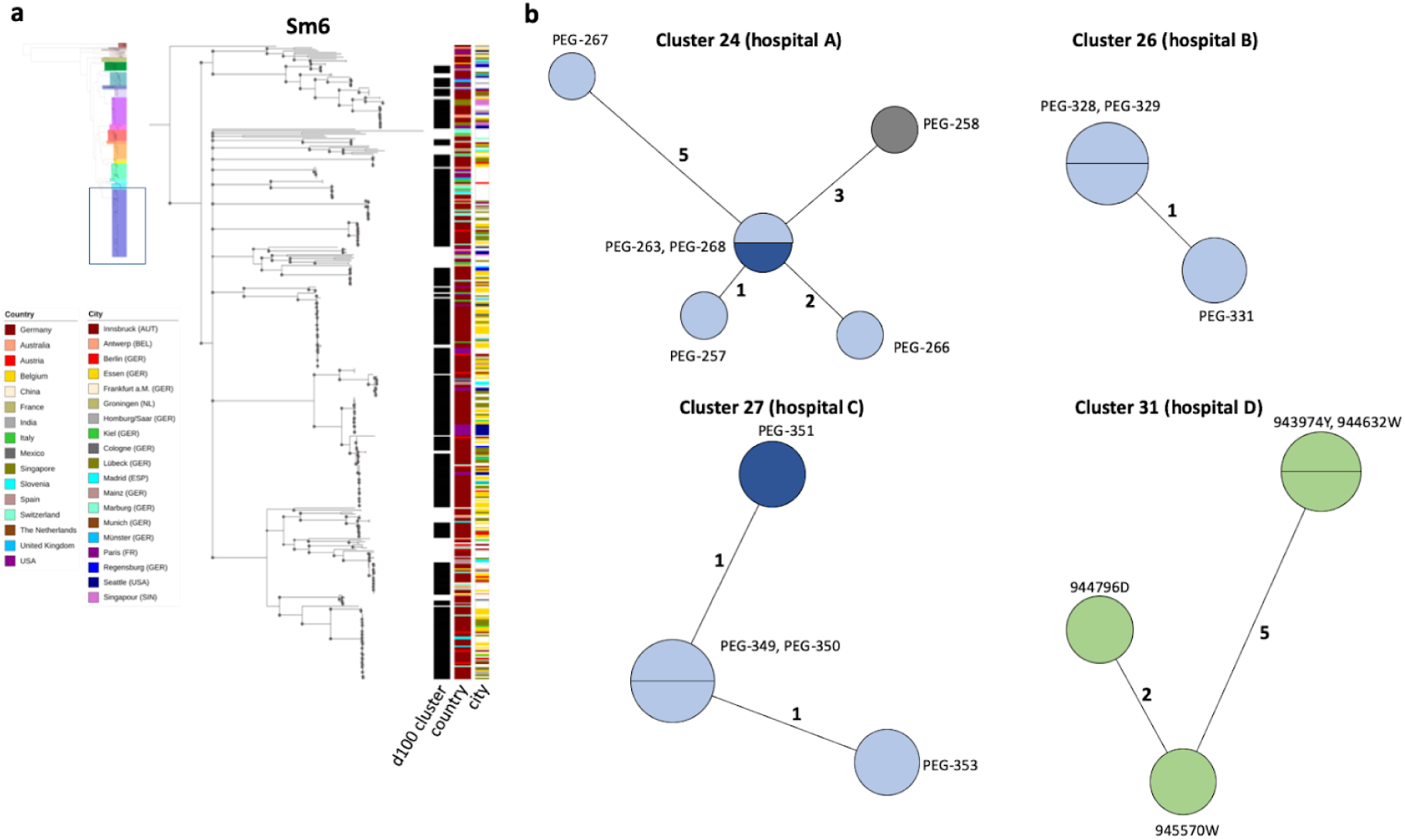
Analysis of d100 clusters in lineage Sm6 and closely related d10 clusters across the study collection. **a,** The d100 clusters in the largest human-associated lineage Sm6 consist of strains from various countries and, for strains from the same country, of various cities. The coloured bars represent, from left to right, the d100 clonal complexes, the country of isolation, and the city of isolation. **b,** High-resolution analysis of four selected d10 allele clusters for which detailed metadata, i.e. day, source, and ward of isolation, was available are shown as Minimum Spanning Trees based on the 100% core genome MLST loci of the respective cluster. The number of loci used were 3734 for cluster 24 (hospital A), 4190 for cluster 26 (hospital B), 3637 for cluster 27 (hospital C), and 3714 for cluster 31 (hospital D). The number of mismatched alleles are shown in small numbers on the connecting branches. Node colours indicate isolation source, light blue = respiratory sample, dark blue = sputum, grey = wound swap, green = endoscope.

Some strains of lineages, notably those with primarily environmental strains, did not cluster at d10 level at all (Sgn1-4, Sm1, Sm15) (Table S5). The d10 clustering rate ranged from 18% for strains of lineage Sm4a to 48% for Sm13, while 21% of lineage Sm6 strains were in d10 clusters. When year and location of isolation were known and included in further investigations of the d10 clusters, we detected a total of 59 strains, grouped into 13 clusters, which were isolated from the same respective hospital in the same year. Of these, three d10 clusters consisted of strains isolated from the respiratory tract of different patients treated in the same hospital within an eight weeks time span or less (Table 1, Fig. 5B, Fig. S6).

## Discussion

The findings of this study demonstrate that strains of the human opportunistic pathogen *S. maltophilia* can be subdivided into 23 monophyletic lineages, with two of these comprising exclusively environmental strains. The remaining lineages contain strains from mixed environmental and human sources. Among these strains, certain lineages such as Sm6 are most frequently found to be human-invasive, human-non-invasive, or human-respiratory strains, pointing towards a potential adaptation to human infection and enhanced virulence. This is further supported by their association with antibiotic resistance genes, resulting in the multidrug resistance observed among human-associated lineages. Our data provide evidence for the global prevalence of particular circulating lineages with hospital-linked clusters collected within short time interval suggesting transmission. The latter emphasizes the need to instate or re-enforce hygiene and infection control practices to minimize in hospital spread of these pathogens.

In line with previous reports, our large genome-based study revealed that the *S. maltophilia* complex is extraordinarily diverse at the nucleotide level, representing a challenge for population-wide analyses and molecular epidemiology^8, 16–18^. To address this, we first developed a new genome-wide gene-by-gene typing scheme, consisting of 17,603 gene targets, or loci. This whole-genome MLST typing scheme provides a versatile tool for genome-based analysis of *S. maltophilia* complex strains and a unified nomenclature to facilitate further research on the complex with an integrative genotyping tool and sequence data analysis approach. Including the loci of the 7-gene classical MLST typing scheme as well as the *gyrB* gene enables backwards compatibility and comparison of allele numbers with sequence types obtained through the classical MLST scheme^25^. Applying the wgMLST approach to our extensive and geographically diverse collection of *S. maltophilia* strains allowed us to infer a comprehensive phylogenetic population structure of the *S. maltophilia* complex, including the discovery of six previously unknown lineages in addition to those described previously^8, 16^.

Altogether, we found 23 distinct phylogenetic lineages of the *S. maltophilia* complex, which are well supported by hierarchical Bayesian clustering analysis of the core genome and intra- and inter-lineage average nucleotide identity. This genetic heterogeneity observed between the detected lineages is sufficient to consider them as clearly separate lineages of the *S. maltophilia* complex, in line with previous results from classical typing methods and phylogenetic studies^8, 16–18, 28^. In parallel with these reports, human adaptation is observed to vary, with strains from lineages Sgn1, Sgn2, Sgn3, and Sm11 mostly isolated from the environment and strains from the other lineages mostly derived from human or human-associated sources. Apart from the purely environmental lineages Sgn1 and Sgn2, our results indicate that strains from all other lineages are able to colonize humans and cause infection, including lineage Sgn4 outside the “*sensu lato*” group, and potentially switch back and forth between surviving in the environment and within a human host. These results do not support the notion that the *S. maltophilia sensu stricto* strains of lineage Sm6 represent the primary human pathogens^16^. We therefore propose to use the term *S. maltophilia* complex and the respective lineage classification for all strains that are identified as *S. maltophilia* by routine microbiological diagnostic procedures in hospitals and omit the use of *sensu stricto* or *lato*.

Our results further illustrate that strains of nearly all 23 lineages are present in sampled countries and continents, suggesting a long evolutionary trajectory of *S. maltophilia*, likely from an exclusively environmental lifestyle towards human colonization and infection. As almost all lineages contain isolates from environmental sources, pathoadaptation of environmental strains to human colonization and pathogenicity likely occurred multiple times, or individual strains emerged that are adapted to survive in both niches. A recent study based on 80 *Legionella* spp. genomes illustrated that the capacity to infect eukaryotic cells has been acquired independently many times within the genus^39^. The evolution within the *S. maltophilia* complex might have been aided by the apparent genomic plasticity as seen from quite distinct genome lengths and structural variation, even within individual lineages. In addition, multiple pathoadaption events could constitute one of the causes for the relatively large accessory genome we detected. A striking observation achieved by long-read PacBio sequencing was the absence of plasmids that hence did not play a role in gene exchange and resistance development in the selected *S. maltophilia* strains.

It is well established that *S. maltophilia* is equipped with an armamentarium of antimicrobial resistance-conferring mechanisms^5, 20^. In our strain collection, we found several families of antibiotic efflux pumps ubiquitously present among strains of all 23 lineages, as well as other genes implicated in aminoglycoside or fluoroquinolone resistance. In some cases, resistance-related genes were only present in some lineages, such as the ß-lactamase gene *blaL2* or the aminoglycoside acetyl- and phosphotransferases genes *aac* and *aph*. A striking finding was that the four lineages most distantly placed from the remaining *S. maltophilia* complex, Sgn1 - Sgn4, were associated with the lack of key virulence and resistance factors. In contrast, the most successful human-associated lineage Sm6 was linked to the presence of ß-lactamases (BlaL1 and BlaL2) and aminoglycoside resistance-conferring enzymes (Aph) as well as KatA, involved in resistance to disinfectants, pointing towards adaptation to healthcare settings and survival on and in patients. This finding might explain why strains of lineage Sm6 were dominant in our investigation, both in our total study collection as well as in the subset of prospectively collected strains as the majority of strains were isolated from human-associated sources. This notion is also supported by our finding that we did not detect any d100 clusters, or circulating variants, in the primarily environmental-associated lineages.

Importantly, our study indicates the presence of potential transmission clusters in human-associated strains, suggesting potential direct or indirect human-to-human transmission^8^. Indeed, we identified a remarkable number of closely related strains (270) that congregated in 62 clusters as indicated by a maximum of ten mismatched alleles in the pairwise comparison. While no d10 clusters were found in the more distantly placed lineages Sgn1-4, all other lineages comprised such clusters with similar clustering rates. A common source of infection is supported in those cases where detailed epidemiological information concerning hospital and day of isolation was available. Further studies looking into potential transmission events are warranted as this would have major consequences on how infection prevention and control teams deal with *S. maltophilia* colonization or infection.

Our study is limited by our collection framework. Molecular surveillance of *S. maltophilia* is currently not routinely performed and no robust data on prevalence, sequence types, or resistance profiles exist. The geographic restriction of our prospective sampling is biased towards the acquisition of clinical and human-pathogenic *S. maltophilia* strains from a multinational consortium mainly comprised of German, Austrian, and Swiss hospitals. The inclusion of all available sequence data in public repositories compensates this restriction partially, however, for these strains information on isolation source and date was incomplete or missing. More prospective, geographically diverse sampling from different habitats is warranted to corroborate our findings, especially concerning the apparent habitat adaptation to the human host. Ultimately, it will be highly interesting to correlate genotype to patient outcomes to identify genomic groups that might be associated with a higher virulence.

Taken together, our data show that strains from several diverse *S. maltophilia* complex lineages are associated with the hospital setting and human-associated infections, with lineage Sm6 strains potentially best adapted to colonize or infection humans. Strains of this lineage are isolated worldwide, are found in potential human-to-human transmission clusters, and are predicted to be highly resistant to antibiotics and disinfectants. Accordingly, strict compliance to infection prevention measures is important to prevent and control nosocomial transmissions especially of *S. maltophilia* lineage Sm6 strains, including the need to ensure that the commonly used disinfectants are effective against *S. maltophilia* complex strains expressing KatA. Future anti-infective treatment strategies may be based on our finding of a very low prevalence (1.7%) of trimethoprim-sulfomethoxazole resistance genes in our collection, suggesting that this antibiotic drug remains the drug of choice for the treatment of *S. maltophilia* complex infections.

## Online methods

### Bacterial strains and DNA isolation

All *Stenotrophomonas maltophilia* complex strains sequenced in this study were routinely collected in the participating hospitals and identified as *S. maltophilia* using MALDI-TOF MS. The strains were grown at 37°C or 30°C in either lysogeny broth (LB) or Brain Heart Infusion media. RNA-free genomic DNA was isolated from 1-ml overnight cultures using the DNeasy Blood & Tissue Kit according to the manufacturer’s instructions (Qiagen, Hilden, Germany). To ensure correct taxonomic identification as *S. maltophilia*, the 16S rRNA sequence of K279a (NC_010943.1) was blasted against all strains. The large majority (1,278 strains, 98%) of our dataset had 16S rRNA similarity values ≥99% (rounded to one decimal). 27 strains, mostly from the more distant clades Sgn1-4, had 16S rRNA blast results between 98.8% and 98.9%. Where no 16S rRNA sequence was found (one study using metagenome assembled genomes^40^ as well as accession numbers GCA_000455625.1 and GCA_000455685.1) we left the isolates in our collection if the allele calls were above the allele threshold of 2,000 (Fig S1H).

### Whole genome data collection and next generation sequencing

We retrieved available *S. maltophilia* sequence read datasets and assembled genomes from NCBI nucleotide databases as of April 2018, excluding next generation sequencing (NGS) data from non-Illumina platforms and datasets from studies that exclusively described mutants. For studies investigating serial strains from the same patient, we chose only representative strains, i.e. one sample per patient was chosen from Esposito *et al.*^41^ and one strain of the main lineages found by Chung *et al.*^42^. In case of studies providing both NGS data and assembled genomes, we included the NGS data in our analysis.

In addition, we sequenced the genomes of 1,071 clinical and environmental strains. NGS libraries were constructed from genomic DNA using a modified Illumina Nextera protocol^43^ and the Illumina NextSeq 500 platform with 2×151bp runs (Illumina, San Diego, CA, United States). NGS data was assembled *de novo* using SPAdes (v3.7.1) included into the BioNumerics software (v7.5, Applied Maths NV). We excluded assemblies with an average coverage depth < 30x (Fig. S1A), deviating genome lengths (< 4Mb and > 6Mb) (Fig. S1B), number of contigs > 500 (Fig. S1C), more than 2000 non-ACTG bases (Fig. S1D), an average quality < 30 (Fig. S1E), and GC content (<63% or >68%) (Fig. S1F). 55 datasets where assembly completely failed were excluded from further analysis. For the phylogenetic analysis, we further excluded strains possessing less than 2,000 genes of the whole genome MLST scheme constructed in this study (Fig. S1H). The resulting dataset contained 1,305 samples (255 from public databases) with a mean coverage depth of 130x (SD = 58; median 122, IQR 92-152), consisted of, on average, 74 contigs (mean, SD = 44; median 67, IQR 47 – 93), and encompassed a mean length of 4.7 million base pairs (SD = 0,19; median 4.76, IQR 4.64 – 4.87) (Data S4). Next generation sequencing data generated in the study is available from public repositories under the study accession number PRJEB32355 (accession numbers for all datasets used are provided in Data S3).

### Generation of full genomes by PacBio third generation sequencing

We used PacBio long-read sequencing on an RSII instrument (Pacific Biosciences, Menlo Park, CA, USA) to generate fully closed reference genome sequences of *S. maltophilia* complex strains sm454, sm-RA9, Sm53, ICU331, SKK55, U5, PEG-141, PEG-42, PEG-173, PEG-68, PEG-305, PEG-390 which together with available full genomes represent the majority of the diversity of our collection. SMRTbellTM template library was prepared according the Procedure & Checklist - 20 kb Template Preparation using the BluePippinTM Size- Selection System (Pacific Biosciences, Menlo Park, CA, USA). Briefly, for preparation of 15-kb libraries, 8 μg of genomic DNA from *S. maltophilia* strains was sheared using g-tubesTM (Covaris, Woburn, MA, USA) according to the manufacturer’s instructions. DNA was end-repaired and ligated overnight to hairpin adapters applying components from the DNA/Polymerase Binding Kit P6 (Pacific Biosciences, Menlo Park, CA, USA). BluePippinTM Size-Selection to 7,000 kb was performed as instructed (Sage Science, Beverly, MA, USA). Conditions for annealing of sequencing primers and binding of polymerase to purified SMRTbellTM template were assessed with the Calculator in RS Remote (Pacific Biosciences, Menlo Park, CA, USA). SMRT sequencing was carried out on the PacBio RSII (Pacific Biosciences, Menlo Park, CA, USA) taking one 240-minutes movie for each SMRT cell. In total 1 SMRT cell for each of the strains was run. For each of the 12 genomes, 59,220 to 106,322 PacBio reads with mean read lengths of 7,678 to 13,952 base pairs (bp) were assembled using the RS_HGAP_Assembly.3 protocol implemented in SMRT Portal version 2.3.0^44^. Subsequently, Illumina reads were mapped onto the assembled sequence contigs using BWA (version 0.7.12)^45^ to improve the sequence quality to 99.9999% consensus accuracy. The assembled reads were subsequently disassembled for removal of low-quality bases. The contigs were then analysed for their synteny to detect overlaps between its start of the anterior and the end of the posterior part to circularise the contigs. Finally, the *dnaA* open reading frame was identified and shifted to the start of the sequence. To evaluate structural variation, genomes were aligned using blastn. PasmidFinder was used to screen the completed genomes for plasmids^46^. Genome sequences are available under bioproject number; the accession numbers can be found in Table S2.

### Construction of a whole-genome MLST scheme for the *S. maltophilia* complex

A whole-genome multilocus sequence typing (wgMLST) scheme was created by Applied Maths NV (bioMérieux) using 171 publically available *S. maltophilia* genome datasets. First, an initial set of loci was determined using the coding sequences (CDS) of the 171 genomes (Data S1). Within this set, loci that overlapped more than 75% or that yielded BLAST hits at the same position within one genome were omitted or merged until only mutually exclusive loci were retained while preserving maximal genome coverage. Mutually exclusive loci are defined as loci for which the reference alleles (typically one or two unique DNA sequences per loci) only yield blast hits at a threshold of 80% similarity to their own genomic location and not to reference alleles of another locus, such as paralogs or repetitive regions. In addition, loci that had a high ratio of invalid allele calls (e.g. because of the absence of a valid start/stop codon [ATG, CTG, TTG, GTG], the presence of an internal stop codon [TAG, TAA, TGA], or non-ACTG bases) and loci for which alleles were found containing large tandem repeat areas were removed. Lastly, multi-copy loci, i.e. repeated loci for which multiple allele calls were retrieved, were eliminated to achieve 90% of the genome datasets used for scheme validation had less than 10 repeated loci. The resulting scheme contained 17,603 loci (including the seven loci from the previously published MLST scheme^25^, see Table S1) (Fig. S2, Fig. S3) and can be accessed through a plugin in the BioNumerics^TM^ Software (Applied Maths NV, bioMérieux). On average 4,174 loci (range 3,024 - 4,536) were identified per genome of our study collection.

To determine the allele number(s) corresponding to a unique allele sequence for each locus present in the genome of a strain, two different algorithms were employed: the assembly-free (AF) allele calling uses a k-mer approach starting from the raw sequence reads while the assembly-based (AB) allele calling performs a lastn search against assembled genomes with the reference alleles of each loci as query sequences. After each round of allele identification, all available data from the two algorithms (AF and AB) were combined into a single set of allele assignments, called consensus calls. If both algorithms returned one or multiple allele calls for a given loci, the consensus is defined as the allele(s) that both analyses have in common. If there is no overlap, there will be no allele number assigned for this particular locus. If for a specific locus the allele call is only available for one algorithm, this allele call will be included. If multiple allele sequences were found for a consensus locus, only the lowest allele number is retained. Only those genes are assigned an allele number that have valid start/stop codons and do not exceed a defined maximum of ambiguous bases and N’s. The loci of the scheme were annotated using the blast2go tool^47^ relying on NCBI blast version 2.4.0+ ^48^ and InterProscan 5 online^49^ (Data S2), and the November 2018 GO^50, 51^ and NCBI nr databases were used.

### Whole genome Multilocus sequence typing scheme validation

To validate the scheme, a collection of repeatedly sequenced ATCC strains^52^ as well as sequence reads sets from published work^41^ were analyzed with the wgMLST scheme in BioNumerics (v7.6.3). For the technical replicates (same sequence read set analysed multiple times), the number of consensus allele calls and allelic profiles were identical. Biological replicates (sequencing data obtained from different fresh cultures of *S. maltophilia* strain ATCC 13637^52^) differed at maximum 5 consensus allele calls.

### Phylogenetic analysis

We characterized the core loci present in 99% of the dataset based on loci presence, i.e. that genes received a valid allele call, amounting to 1,275 loci. For phylogenetic analyses, a concatenated alignment of the 1,275 core genes from all strains was created, and an initial tree was built using RAxML-NG with a GTR+Gamma model and using the site-repeat optimization^53^. This alignment and tree was then fed to ClonalFrameML to detect any regions of recombination^54^. These regions were then masked using maskrc-svg and this masked alignment was then used to build a recombination-free phylogeny using the same approach as above in RAxML-NG. iTOL was employed for annotating the tree^55^.

We detected phylogenetic lineages within the tree using a hierarchical Bayesian Analysis of Population Structure (hierBAPS) model as implemented in R (rHierBAPs) with a maximum depth of 2 and maximum population number of 100^56^. FastANI^57^ was employed to calculate the pairwise Average Nucleotide Identity (ANI) as a similarity matrix between all the strains with the option ‘many-to-many’. The similarity matrix was imported into R and used together with the group assignment obtained from hierBAPS to compare the ANI values in strains within and between groups. ANI values were plotted as a heatmap of all strains as well as a composite histogram of identity between and within groups.

### Identification of lineage-specific loci

To detect lineage- and isolation source-specific loci, the allele database obtained from wgMLST analysis in BioNumerics for the 1,305 *S. maltophilia* strains was filtered using Base R version 3.4.3^58^ and the tidyverse package^59^ to identify loci that were uniquely present in the phylogenetic lineages or specific for isolation source. Intersecting gene sets were visualized with UpSetR^60^.

### Resistome and virulence analysis

Resistome and virulome were characterized with abricate version 0.8.7^37^ screened against the NCBI Bacterial Antimicrobial Resistance Reference Gene Database (NCBI BARRGD, PRJNA3134047) and the Virulence Factors of Pathogenic Bacteria Database (VFDB)^38^. All genes below 90% coverage breadth were excluded. In addition, literature was reviewed to identify additional genes associated with antibiotic resistance and virulence in *S. maltophilia*.

### Statistical analysis and data management

All statistical analyses and data management were performed in R version 3.4.3^58^ using mainly packages included in the tidyverse^59^, reshape2^61^, and rcompanion^62^. The map was created with the tmap package^63^. WgMLST data was analyzed in BioNumerics v7.6.3 using the WGS and MLST plugins. For correlation testing between variables Spearman’s rank test was employed. The test of equal or given proportions or Fisher’s exact test for sample sizes smaller than 5 was used to test for proportions. Multiple correspondence analysis (MCA) was performed with FactoMineR^64^ and its results were visualised with the factoextra^65^ R packages.

## Supporting information

Genomes used for wgMLST scheme creation

wgMLST loci details

Isolate collection metadata

Sequence metrics

## Acknowledgments

We thank V. Mohr, F. Boysen, and T. Ubben for technical assistance during next generation sequencing of strains. Parts of the work have been funded by grants from German Center for Infection Research, Federal Ministry of Education and Research, Germany, from Deutsche Forschungsgemeinschaft (DFG, German Research Foundation) under Germany’s Excellence Strategy – EXC 22167-390884018, and grants from the Leibniz Science Campus EvoLUNG; IA and WRS received funds from the University of Hamburg; CFL and JWAR were funded through the European Commission Horizon 2020 Framework Marie Skłodowska-Curie Actions (Grant agreement number: 713660 - PRONKJEWAIL - H2020-MSCA-COFUND-2015); CJM was funded by the Department of Economy, Science and Innovation of the Flemish Government and the Belgian Science Policy (Belspo); KZ was funded by the National Natural Science Foundation of China (grant number 81702045), and 13th Five-Year National Major Science and Technology Projects of China (grant number 2018ZX10712001); OCS, DY, IG and XD were funded by the Spanish Ministry for Science, Innovation and Universities (grant reference BIO2015-66674-R); NAR was funded by the SingHealth DUKE-NUS Pathology Academic Clinical Programme Clinical Innovation Support Grant (09/FY2017/P1/06-A20); UN was funded by the EU Horizon 2020 programme, grant agreement number 643476.

## Author contributions

MIG, JR, JS, SN, and TAK conceived the study. MIG, IB, MG, OCS, JS, TAK curated the data. MIG, CJM, IB, MD, AG, MS, ICS, CU, CFL, DY, MRF, OCS and TAK performed the formal analysis. JR, JS, and SN acquired funding. SK, NAR, MK, WRS, KDB, KZ, TS, JWAR, UN, DY, IG, XD, JR, UM, MRF and JS provided resources for this study. MIG, IB, AG, and CFL visualized the results. MIG, JS, SN, TAK wrote the initial draft. All authors critically reviewed and modified the manuscript.

## Data and materials availability

Sequencing data are available under study accession numbers PRJEB32355, PRJEB32585, and PRJNA543082.

## Supplementary Materials

Figures S1 - S6

Tables S1 - S5

Data S1 - S4

