## Supplemental figures and tables for "The global phylogenetic landscape and nosocomial spread of the multidrug-resistant opportunist *Stenotrophomonas maltophilia*"

**This PDF file includes:**

Figs. S1 to S6
Tables S1 to S5
Captions for Data S1 to S4

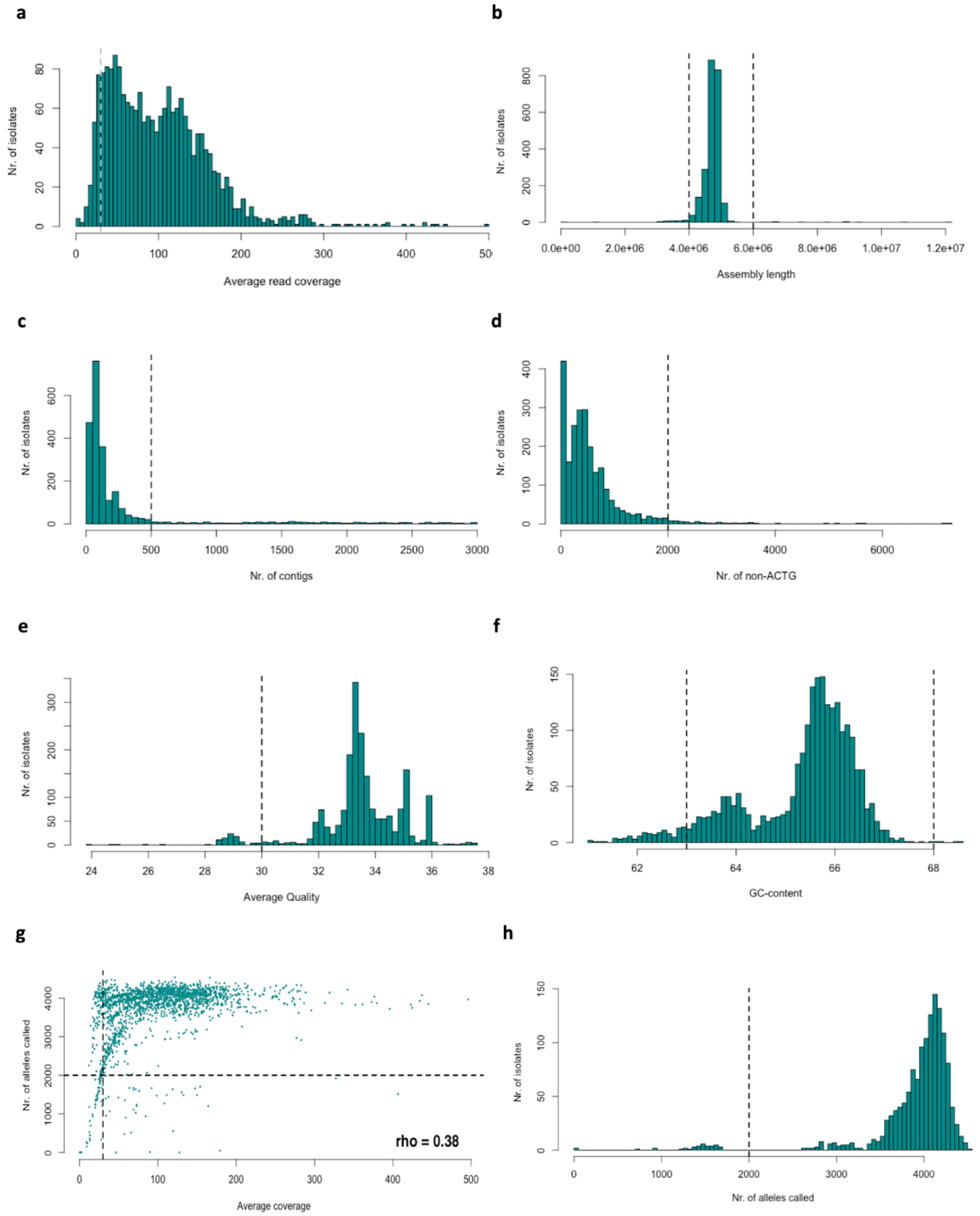


**Fig. S1. Quality metrics upon assembly of a collection of 2,389 *S. maltophilia* complex genome datasets before filtering**. Dashed lines indicate the quality thresholds applied in this study. Barplots. **a,** average read coverage. **b,** genome lengths upon assembly. **c,** number of contigs. **d,** number of non-ACTG bases called. **e,** average quality. **f,** GC-content. **g,** Scatterplot of the number of allele calls received by the isolates versus coverage

(Spearman’s rank correlation coefficient shown on the right lower side of the figure). **h,** number of loci that were called and received an allele number of the wgMLST scheme.

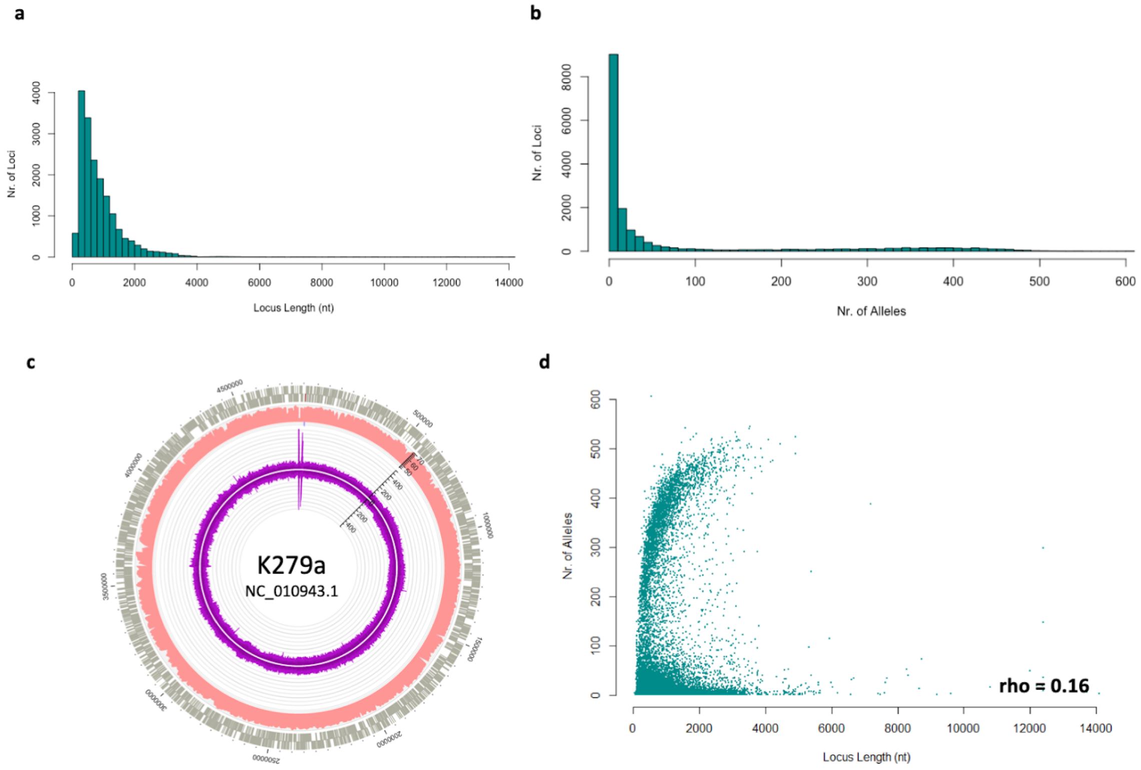


**Fig. S2. Characteristics of the 17,603 loci of the wgMLST scheme constructed for the *S. maltophilia* complex**. **a,** Distribution of the wgMLST loci lengths. **b,** Distribution of the number of different alleles per locus. **c,** Location of the wgMLST loci that map to the clinical reference strain *S. maltophilia* K279a. Coverage is depicted in purple, GC content in red. Impact of locus length on allele diversity (n = 1,305 genomes). **d,** Scatter plot of number of alleles versus locus length (Spearman’s rank correlation coefficient shown on the right lower side of the figure).

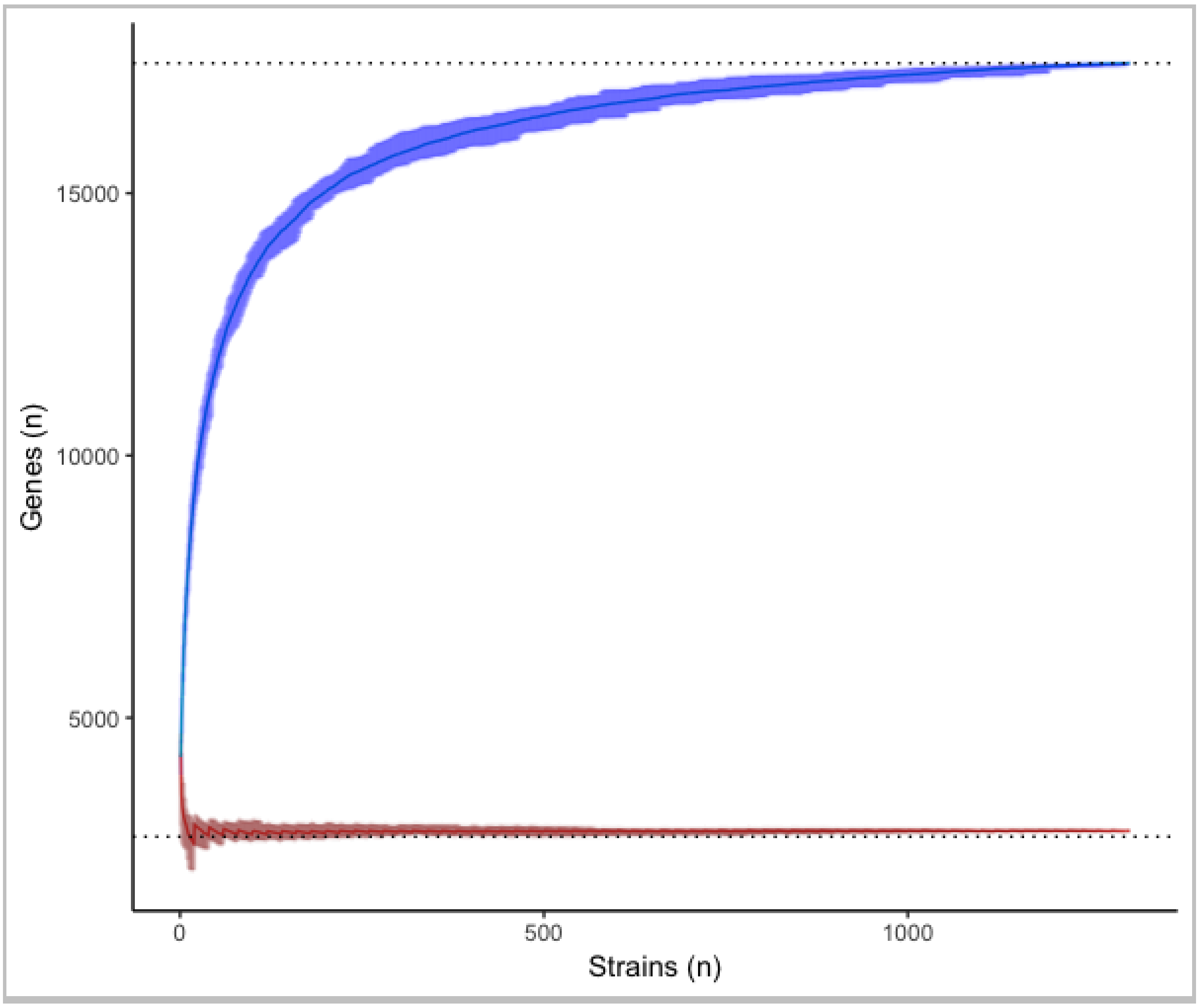


**Fig. S3. Rarefaction curve of the core- and pangenome of the 1,305 *S. maltophilia* complex strains in the study collection.** Analysis based on the presence of genes in the assemblies and independent of the number of received allele calls for the respective loci of the wgMLST scheme. The x axis shows the number of strains taken into the analysis and the y axis displays the number of genes detected in the assembled genomes of these strains. Upon 100 repeats of random selection of genomes from the complete set, the minimum and maximum of each calculation are shown in shaded colours with the average as solid line. Blue line represents the pan genome, red line the 95% core genome.

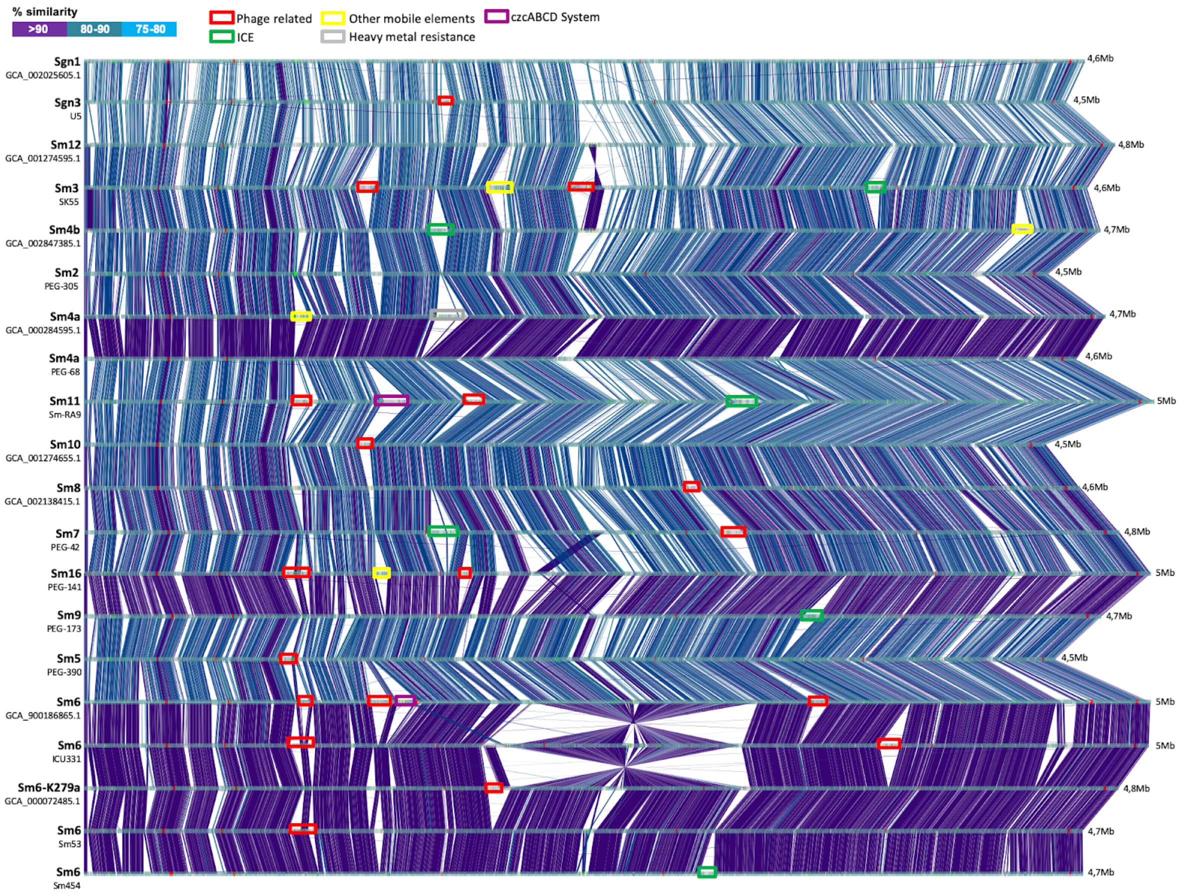


**Fig. S4**. **Structural variation indicated by a genome-wide alignment of 20 *S. maltophilia* complex genomes using blastn**. These genomes are representative of the 15 major phylogenetic lineages and include the clinical reference strain K279a (accession NC_010943.1). The alignment shows major structural variation and different genome lengths of strains from different lineages and even for strains from the same lineage for lineages Sm4a and Sm6. One strain, ICU331, exhibited a large inversion of ~1Mb, as verified upon aligning the reads on the assembly and flanked by several IS elements. Colored squares show genomic islands found across the genomes. ICE = Integrative and conjugative elements.

**a**

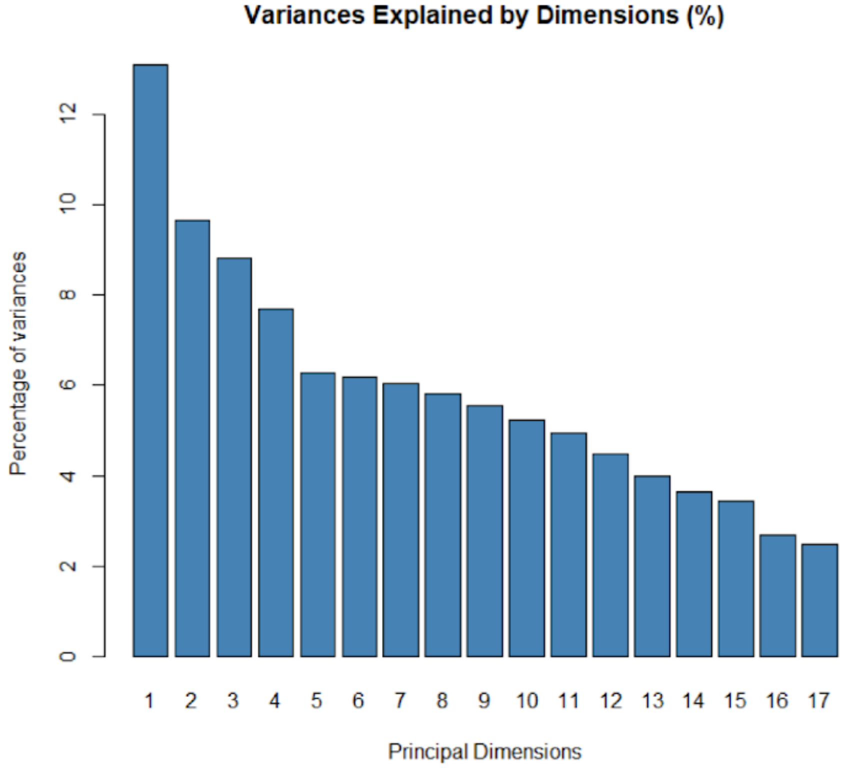
Variances Explained by Dimensions (%)
Percentage of variances
0
2
4
6
8
10
12
1 2 3 4 5 6 7 8 9 10 11 12 13 14 15 16 17
Principal Dimensions

**b**

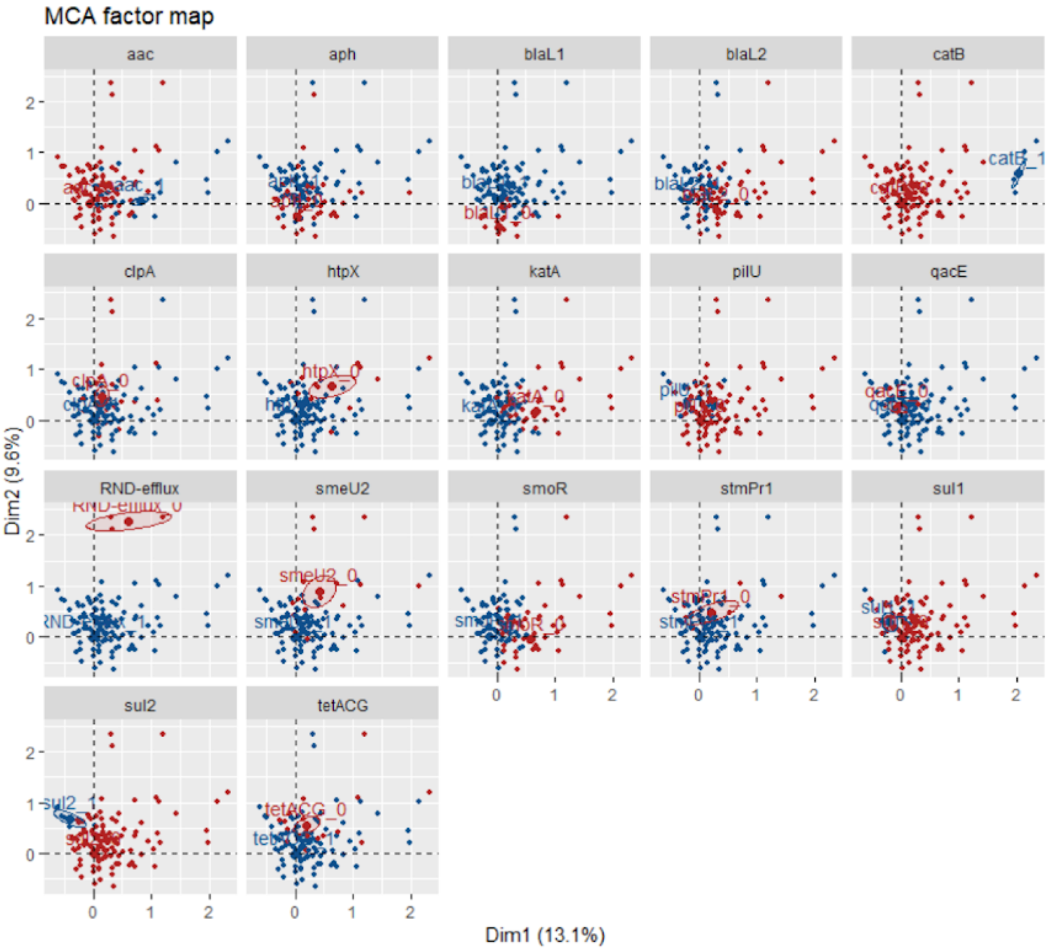
MCA factor map
aac
aph
blaL1
blaL2
catB
clpA
htpX
katA
pilU
qacE
RND-efflux
smeU2
smoR
stmPr1
sul1
sul2
tetACG
catB_1
htpX_0
smeU2_0
stmPr1_0
tetACG_0
sul2_1
Dim2 (9.6%)
Dim1 (13.1%)
0
1
2

**Fig. S5. Multiple correspondence analysis of 17 resistance or virulence genes.** Barplot displaying the percentage of variance explained by the respective principal component dimensions of the multiple correspondence analysis (MCA). First four dimensions give us the percentage of variance explained by the model while the rest are individual variations of active variables. **a,** MCA results shown on individual basis and grouped by each of the 17 resistance or virulence associated genes or groups of genes (RND-efflux) acting as active variables. **b,** Presence/absence of a gene within an individual denoted with blue and red points respectively.

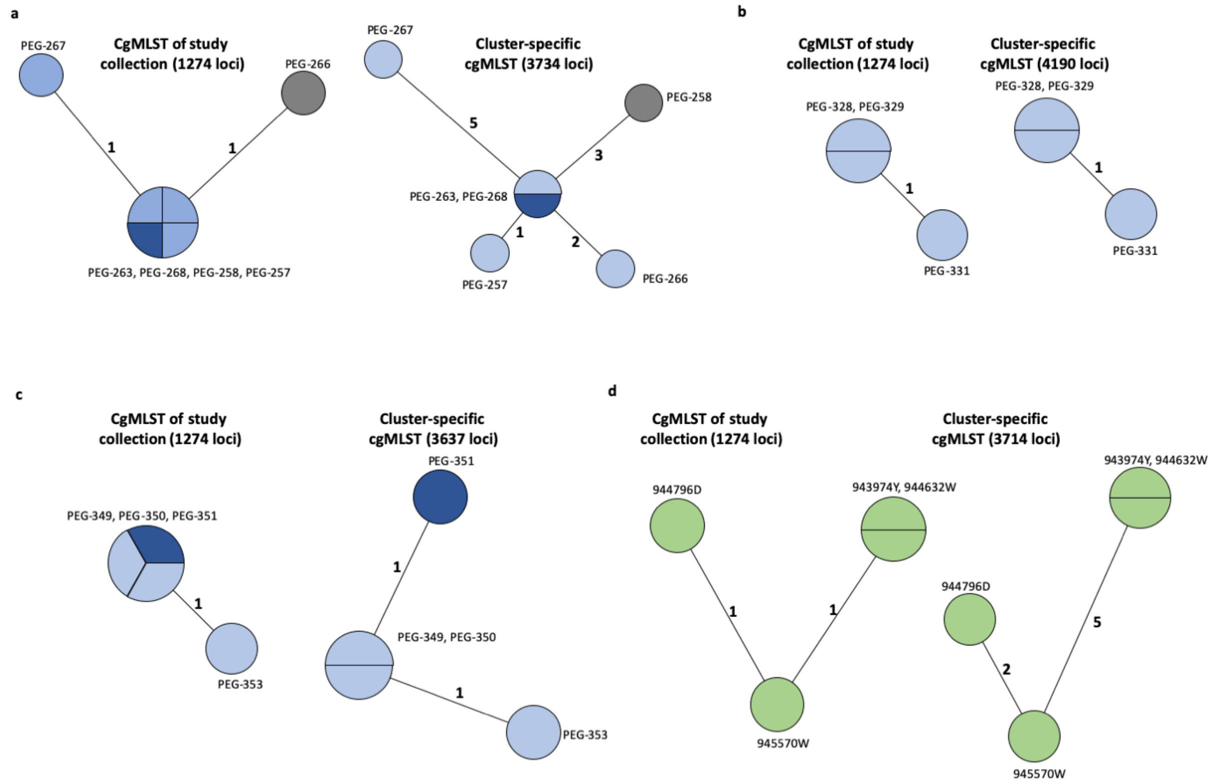


**Fig. S6. Minimum Spanning Trees based on either the 1,274 study collection core genome MLST loci or the cluster specific core genome MLST loci as indicated above the respective trees**. **a,** Cluster 24 (hospital A). **b,** Cluster 26 (hospital B). **c,** Cluster 27 (hospital C). **d,** Cluster 31 (hospital D). The number of mismatched alleles are shown in small numbers on the connecting branches. Node colours indicate isolation source, light blue = respiratory sample, dark blue = sputum, grey = wound swap, green = endoscope.

**Table S1.**

Details of the 7 classical Multilocus Sequence Typing loci (available at https://pubmlst.org/smaltophilia/ [accessed February 10th, 2019])

| Gene name | Length | Min length (bp) | Max length (bp) | wgMLST loci (MLST sequence) | wgMLST loci (full sequence) |
|---|---|---|---|---|---|
| *atpD* | Fixed | 531 | 531 | STENO_17597 | STENO_3197 |
| *gapA* | Variable | 558 | 561 | STENO_17598 | STENO_2953 |
| *guaA* | Fixed | 552 | 552 | STENO_17599 | STENO_1653 |
| *mutM* | Fixed | 465 | 465 | STENO_17600 | STENO_47 |
| *nuoD* | Fixed | 444 | 444 | STENO_17601 | STENO_2646 |
| *ppsA* | Variable | 492 | 495 | STENO_17602 | STENO_2281 |
| *recA* | Fixed | 546 | 546 | STENO_17603 | STENO_1423 |

**Table S2.**

Accession numbers, isolation source, and sequence metrics of 20 fully finished genomes from the major lineages, 12 of which have been sequenced within this study.

| Isolate name | Accession | Lineage | Sequenced | Source | Mean read length (bp) | Mean coverage | No. of contigs | Length (Mb) | No. of genes | Plasmids |
|---|---|---|---|---|---|---|---|---|---|---|
| sm454 | CP040431 | Sm6 | this study | human | 12822 | 200 | 1 | 4,68 | 4210 | 0 |
| ICU331 | CP040440 | Sm6 | this study | human | 9119 | 135 | 1 | 4,90 | 4563 | 0 |
| SKK55 | CP040433 | Sm3 | this study | human | 11162 | 142 | 1 | 4,67 | 4206 | 0 |
| PEG-141 | CP040439 | Sm16 | this study | human | 12112 | 116 | 1 | 5,00 | 4605 | 0 |
| PEG-42 | CP040435 | Sm7 | this study | human | 12863 | 154 | 2 | 4,87 | 4427 | 0 |
| PEG-173 | CP040438 | Sm9 | this study | human | 10384 | 144 | 1 | 4,76 | 4238 | 0 |
| PEG-68 | CP040434 | Sm9 | this study | human | 8830 | 150 | 1 | 4,64 | 4143 | 0 |
| PEG-305 | CP040437 | Sm2 | this study | human | 13952 | 165 | 1 | 4,50 | 3987 | 0 |
| U5 | CP040429 | Sgn3 | this study | environmental | 13269 | 125 | 3 | 4,50 | 4064 | 0 |
| PEG-390 | CP040436 | Sm5 | this study | human | 12253 | 183 | 1 | 4,55 | 4119 | 0 |
| sm-RA9 | CP040432 | Sm11 | this study | environmental | 12190 | 132 | 1 | 5,00 | 4471 | 0 |
| Sm53 | CP040430 | Sm6 | this study | human | 7678 | 119 | 1 | 4,68 | 4199 | 0 |
| GCA_002025605.1 | | Sgn1 | NCBI | environmental | | | 1 | 4,67 | | 0 |
| GCA_001274595.1 | | Sm12 | NCBI | human | | | 1 | 4,8 | | 0 |
| GCA_002847385.1 | | Sm4b | NCBI | unknown | | | 1 | 4,74 | | 0 |
| GCA_000284595.1 | | Sm4a | NCBI | unknown | | | 1 | 4,77 | | 0 |
| GCA_001274655.1 | | Sm10 | NCBI | human | | | 1 | 4,51 | | 0 |
| GCA_002138415.1 | | Sm8 | NCBI | unknown | | | 1 | 4,67 | | 0 |
| GCA_900186865.1 | | Sm6 | NCBI | unknown | | | 1 | 5 | | 0 |
| GCA_000072485.1 | NC_010943.1 | Sm6 | NCBI | human | | | 1 | 4,85 | 4522 | 0 |

**Table S3.**

Frequencies of strains per lineage and their association with isolation source.

| | Anthropogenic | | Environmental | | Human-invasive | | Human-non-invasive | | Human-respiratory | | total per group |
|---|---|---|---|---|---|---|---|---|---|---|---|
| | n | p-value | n | p-value | n | p-value | n | p-value | n | p-value | |
| Sgn1 | 0 | 0.62 | 12 | <.001 | 0 | 0.50 | 0 | 0.08 | 0 | 0.01 | 12 |
| Sgn2 | 0 | 0.82 | 5 | <.001 | 0 | 0.60 | 0 | 0.47 | 0 | 0.24 | 5 |
| Sgn3 | 2 | 0.58 | 28 | <.001 | 0 | 0.08 | 3 | 0.02 | 4 | <.001 | 37 |
| Sgn4 | 1 | 0.74 | 5 | 0.29 | 3 | 0.59 | 9 | 0.52 | 6 | 0.20 | 24 |
| Sm1 | 0 | 0.91 | 0 | 0.83 | 1 | 0.49 | 1 | 0.59 | 0 | 0.52 | 2 |
| Sm10 | 1 | 0.50 | 5 | 0.54 | 8 | 0.55 | 18 | 0.58 | 30 | 0.52 | 62 |
| Sm11 | 7 | <.001 | 11 | <.001 | 3 | 0.62 | 5 | 0.34 | 3 | <.001 | 29 |
| Sm12 | 9 | <.001 | 3 | 0.52 | 5 | 0.59 | 13 | 0.54 | 18 | 0.47 | 48 |
| Sm13 | 0 | 0.54 | 0 | 0.34 | 2 | 0.59 | 4 | 0.46 | 17 | 0.03 | 23 |
| Sm14 | 1 | 0.52 | 2 | 0.47 | 1 | 0.65 | 2 | 0.60 | 2 | 0.50 | 8 |
| Sm15 | 2 | 0.47 | 0 | 0.47 | 1 | 0.54 | 6 | 0.58 | 9 | 0.54 | 18 |
| Sm16 | 1 | 0.52 | 0 | 0.56 | 1 | 0.65 | 1 | 0.51 | 5 | 0.51 | 8 |
| Sm17 | 0 | 0.60 | 0 | 0.51 | 3 | 0.47 | 6 | 0.47 | 4 | 0.50 | 13 |
| Sm18 | 3 | 0.47 | 1 | 0.47 | 3 | 0.58 | 9 | 0.56 | 17 | 0.51 | 33 |
| Sm2 | 0 | 0.47 | 1 | 0.33 | 4 | 0.59 | 9 | 0.47 | 26 | 0.04 | 40 |
| Sm3 | 2 | 0.54 | 5 | 0.52 | 9 | 0.54 | 23 | 0.52 | 30 | 0.58 | 69 |
| Sm4a | 6 | 0.58 | 4 | <.001 | 21 | 0.47 | 65 | <.001 | 57 | 0.17 | 153 |
| Sm4b | 0 | 0.54 | 1 | 0.54 | 7 | 0.02 | 5 | 0.54 | 8 | 0.54 | 21 |
| Sm5 | 1 | 0.60 | 5 | 0.06 | 3 | 0.52 | 4 | 0.54 | 4 | 0.28 | 17 |
| Sm6 | 8 | 0.06 | 19 | <.001 | 42 | 0.58 | 107 | 0.54 | 190 | <.001 | 366 |
| Sm7 | 2 | 0.52 | 3 | 0.17 | 7 | 0.52 | 29 | 0.47 | 40 | 0.49 | 81 |
| Sm8 | 0 | 0.54 | 0 | 0.44 | 0 | 0.34 | 12 | 0.03 | 8 | 0.56 | 20 |
| Sm9 | 6 | 0.47 | 6 | 0.50 | 8 | 0.54 | 21 | 0.47 | 45 | 0.33 | 86 |
| ungrouped | 0 | 0.84 | 1 | 0.54 | 1 | 0.54 | 1 | 0.68 | 1 | 0.54 | 4 |
| Total per origin | 52 | | 117 | | 133 | | 353 | | 524 | | 1179 |

Note: Either the test of equal or given proportions or, for small sample sizes (n < 5), Fisher's exact test was used in 124 comparisons while controlling the false discovery rate of multiple testing using the Benjamini-Hochberg procedure. * < .05; ** < .01; *** < .001

**Table S4.**

Unique whole genome multilocus sequence typing loci per phylogenetic lineage and isolation source.

| Lineage | n | Clinical (%) | | | | | Environmental (%) | Unknown (%) | Unique loci | | |
|---|---|---|---|---|---|---|---|---|---|---|---|
| | | Human-invasive | Human-non-invasive | Human-respiratory | Human-sputum | Anthropognic | | | 90% | 80% | total |
| Sgn1 | 14 | | | | | | 12 (85.7) | 2 (14.3) | 0 | 0 | 0 |
| Sgn2 | 6 | | | | | | 5 (83.3) | 1 (16.7) | 0 | 0 | 1 |
| Sgn3 | 38 | | 3 (7.9) | 2 (5.3) | 2 (5.3) | 2 (5.3) | 28 (73.7) | 1 (2.6) | 133 | 197 | 727 |
| Sgn4 | 28 | 3 (10.7) | 9 (32.1) | 2 (7.1) | 4 (14.3) | 1 (3.6) | 5 (17.9) | 4 (14.3) | 0 | 0 | 0 |
| Sm1 | 2 | 1 (50) | 1 (50) | | | | | | 0 | 0 | 0 |
| Sm2 | 49 | 4 (8.2) | 9 (18.4) | 13 (26.5) | 13 (26.5) | | 1 (2) | 9 (18.4) | 4 | 6 | 167 |
| Sm3 | 81 | 9 (11.1) | 23 (28.4) | 17 (21) | 13 (16) | 2 (2.5) | 5 (6.2) | 12 (14.8) | 0 | 0 | 257 |
| Sm4a | 164 | 21 (12.8) | 65 (39.6) | 46 (28) | 11 (6.7) | 6 (3.7) | 4 (2.4) | 11 (6.7) | 1 | 1 | 246 |
| Sm4b | 22 | 7 (31.8) | 5 (22.7) | 7 (31.8) | 1 (4.5) | | 1 (4.5) | 1 (4.5) | 0 | 0 | 41 |
| Sm5 | 18 | 3 (16.7) | 4 (22.2) | 3 (16.7) | 1 (5.6) | 1 (5.6) | 5 (27.8) | 1 (5.6) | 2 | 2 | 49 |
| Sm6 | 413 | 42 (10.2) | 107 (25.9) | 90 (21.8) | 100 (24.2) | 8 (1.9) | 19 (4.6) | 47 (11.4) | 2 | 5 | 963 |
| Sm7 | 90 | 7 (7.8) | 29 (32.2) | 32 (35.6) | 8 (8.9) | 2 (2.2) | 3 (3.3) | 9 (10) | 10 | 11 | 122 |
| Sm8 | 21 | | 12 (57.1) | 6 (28.6) | 2 (9.5) | 6 (6.6) | 6 (6.6) | 1 (4.8) | 2 | 2 | 60 |
| Sm9 | 91 | 8 (8.8) | 21 (23.1) | 33 (36.3) | 12 (13.2) | 6 (6.6) | 6 (6.6) | 5 (5.5) | 2 | 2 | 226 |
| Sm10 | 67 | 8 (11.9) | 18 (26.9) | 18 (26.9) | 12 (17.9) | 1 (1.5) | 5 (7.5) | 4 (6) | 0 | 0 | 194 |
| Sm11 | 32 | 3 (9.4) | 5 (15.6) | 1 (3.1) | 2 (6.2) | 7 (21.9) | 11 (34.4) | 3 (9.3) | 8 | 8 | 92 |
| Sm12 | 53 | 5 (9.4) | 13 (24.5) | 10 (18.9) | 8 (15.1) | 9 (17) | 3 (5.7) | 5 (9.4) | 19 | 24 | 99 |
| Sm13 | 25 | 2 (8) | 4 (16) | 13 (52) | 4 (16) | | | 2 (8) | 0 | 0 | 0 |
| Sm14 | 17 | 1 ( (11.1) | 2 (22.2) | 1 (11.1) | 1 (11.1) | 1 (11.1) | 2 (22.2) | 1 (11.1) | 0 | 0 | 0 |
| Sm15 | 19 | 1 (5.3) | 6 (31.6) | 3 (15.8) | 6 (31.6) | 2 (10.5) | | 1 (5.3) | 0 | 0 | 2 |
| Sm16 | 9 | 1 (11.1) | 1 (11.1) | 2 (22.2) | 3 (33.3) | 1 (11.1) | | 1 (11.1) | 0 | 0 | 0 |
| Sm17 | 14 | 3 (21.4) | 6 (42.9) | 2 (14.3) | 2 (14.3) | | | 1 (7.1) | 7 | 7 | 51 |
| Sm18 | 36 | 3 (8.3) | 9 (25) | 15 (41.7) | 2 (5.6) | 3 (8.3) | 1 (2.8) | 3 (8.3) | 12 | 17 | 48 |
| ungrouped | | 1 | 1 | | 1 | | 1 | | | | |
| Total | | 133 | 353 | 316 | 208 | 52 | 117 | 126 | | | |

**Table S5.**

Clustering characteristics of the 1,305 *S. maltophilia* complex strains of the study collection divided by lineage based on 100 (d100 clusters) or 10 allelic mismatches (d10 clusters).

| Lineage (n) | d100 clustering rate | d10 clustering rate | No. of d10 clusters | No. of d10 clustered isolates | No. of isolates in individual d10 cluster |
|---|---|---|---|---|---|
| Sgn1 (14) | 0 | 0 | - | - | - |
| Sgn2 (6) | 0 | 0 | - | - | - |
| Sgn3 (38) | 0 | 0 | - | - | - |
| Sgn4 (28) | 0 | 0 | - | - | - |
| Sm1 (2) | 0 | 0 | - | - | - |
| Sm2 (49) | 0.59 | 0.31 | 3 | 15 | 3 - 8 |
| Sm3 (81) | 0.28 | 0.21 | 4 | 17 | 3 - 6 |
| Sm4a (164) | 0.83 | 0.18 | 8 | 30 | 3 - 4 |
| Sm4b (22) | 0.32 | 0.27 | 1 | 6 | 6 |
| Sm5 (18) | 0.5 | 0.38 | 2 | 7 | 3 - 4 |
| Sm6 (413) | 0.73 | 0.21 | 22 | 88 | 3 - 9 |
| Sm7 (90) | 0.92 | 0.27 | 4 | 24 | 3 - 12 |
| Sm8 (21) | 0.71 | 0.38 | 1 | 8 | 8 |
| Sm9 (91) | 0.42 | 0.21 | 4 | 19 | 3 - 10 |
| Sm10 (67) | 0.19 | 0.18 | 2 | 12 | 3 - 9 |
| Sm11 (32) | 0.22 | 0.22 | 2 | 7 | 3 - 4 |
| Sm12 (53) | 0.36 | 0.08 | 1 | 4 | 4 |
| Sm13 (25) | 0.88 | 0.48 | 2 | 12 | 6 |
| Sm14 (9) | 0.66 | 0.44 | 1 | 4 | 4 |
| Sm15 (19) | 0.74 | 0 | - | - | - |
| Sm16 (9) | 0.88 | 0.44 | 1 | 4 | 4 |
| Sm17 (14) | 0.64 | 0.29 | 1 | 4 | 4 |
| Sm18 (36) | 0.75 | 0.25 | 3 | 9 | 3 |

**Data S1. (separate file)**

List of NCBI GeneBank accession numbers of the 171 assembled genomes used to create the whole-genome Multilocus sequence typing scheme.

**Data S2. (separate file)**

Characteristics of the 17,603 loci of the whole-genome Multilocus Sequence Typing scheme.

**Data S3. (separate file)**

Accession numbers and associated metadata for all 1,305 *S. maltophilia* isolates included in the phylogenetic analysis.

**Data S4. (separate file)**

Sequence metrics, number of consensus whole genome Multilocus sequence typing allele calls of all 1305 *S. maltophilia* isolates and other *Stenotrophomonas* species.
